# Representing experience over time: sustained sensory patterns and transient frontroparietal patterns

**DOI:** 10.1101/2022.08.02.502469

**Authors:** Gal Vishne, Edden M. Gerber, Robert T. Knight, Leon Y. Deouell

## Abstract

Instances of sustained stationary sensory input are ubiquitous. However, previous work focused almost exclusively on the transient onset responses. This presents a critical challenge for neural theories of consciousness, which should account for the full temporal extent of experience. Here we use intracranial recordings from human patients viewing diverse images in multiple durations. In sensory regions, we reveal that despite dramatic changes in activation magnitude, the distributed representation of categories and exemplars remains sustained and stable. In contrast, in frontoparietal regions we find transient content representation at stimulus onset. Our results highlight the connection between the anatomical and temporal correlates of experience. To the extent perception is sustained, it may rely on sensory representations, and to the extent it is discrete, centered on perceptual updating, it may rely on frontoparietal representations.

## Introduction

In essence, every perception has non-zero duration – we gaze at a tree for some time, then shift our gaze to look at a fly which just landed on the table, only to take off after a few seconds. All these experiences have a content (a tree, a fly) which extends not only in space but also in time. Most discussions of the Neural Correlates of Consciousness (NCC), defined as the minimal set of mechanisms that are together necessary and sufficient for any one specific experience^1^, addressed the anatomical location in the brain which gives rise to the experience, while time has received considerably less attention in the NCC literature. Introspectively, it seems that our experience unfolds continuously, in parallel with the sequence of events, thus, we would expect that our experience of gazing at the tree was longer than the quick glance at the fly. However, this intuition is complicated by the existence of postdictive effects, when a current stimulus influences the experience of *prior* events^2,3^. To account for this, some have argued that we are not continuously conscious, but rather, we are conscious at discrete moments in time^4,5^. The anatomical component of the NCC is also debated. One major point of contention involves the role of the prefrontal cortex compared to high-level sensory cortices^6,7^. Multiple previous studies found prefrontal responses to be associated with stimulus awareness, yet more recently, it was argued that this is a byproduct of the reporting procedure, and not a signal pertaining to awareness per-se^8^.

To date, the search for the anatomy of the NCC (where) and the temporal progression (when) of consciousness have progressed largely in parallel, as most studies focused on the transient onset or change-related responses, without examining the ubiquitous periods of stationarity between the changes^9–11^. This is a critical challenge, as theories linking conscious experience to the brain should be able to account for experience of sustained events, not only that of stimulus onsets or changes. Importantly, the few studies which examined the full temporal dynamics of responses to longer stimuli found that activity in high-order visual regions drops dramatically shortly after the initial onset response, independent of stimulus duration^12,13^, posing a problem for theories emphasizing the role of neural ignition which argue for continuous experience^14,15^. However, these studies focused on activation dynamics, and did not examine content representation over time, which is the focus of the current study. Additionally, they focused exclusively on visual regions, without addressing activation or representation in frontoparietal cortex. Testable predictions regarding these issues were recently put forward in an adversarial collaboration aiming to adjudicate between the two theories^16^, Global Neuronal Workspace Theory (GNWT)^17^ and Integrated Information Theory (IIT)^18^.

Here, we address these fundamental questions by examining the spatiotemporal neural representation of clearly visible images of different durations (300-1500ms) in ten human patients with drug-resistant epilepsy implanted with subdural electrodes for clinical purposes (Figure 1A, Tables S1-S3). To maintain attention, responses were required for 10% of the trials which were not analyzed, excluding any report related signals (Figure 1B). Using multiple presentation durations allows us to distinguish between responses to the onset of a stimulus and signals tracking the ongoing stimulus presence. Presenting a diverse set of images enables us to identify signals tracking the content of experience, not only whether an image was shown or not (which was the focus of our previous work using this dataset^12^). We use this to examine visual representation over time at the level of visual categories (e.g., faces\objects) and at the level of single exemplars (e.g., specific objects). Within the object category, we analyzed separately a sub-category of watches, which bear low level visual similarity to faces^19,20^. To foreshadow our results, we find that despite considerable variability in moment-to-moment activation levels, the distributed population representation of visual content (category and exemplar-level) in sensory regions is stable, paralleling the stimulus presentation. Further, we find transient visual representation in prefrontal cortex, despite the lack of overt report. These findings confirm predictions of both theories delineated in the recent adversarial collaboration^16^. More broadly, these results highlight the importance of addressing the NCC within a temporal context: to the extent conscious experience is continuous, it may rely on sensory representations, and to the extent it is discrete, it may rely on prefrontal representations. Thus, by studying visual experience and representation beyond the onset, we reveal a deep connection between spatial (anatomical) and temporal understanding of consciousness.

**Figure 1.**
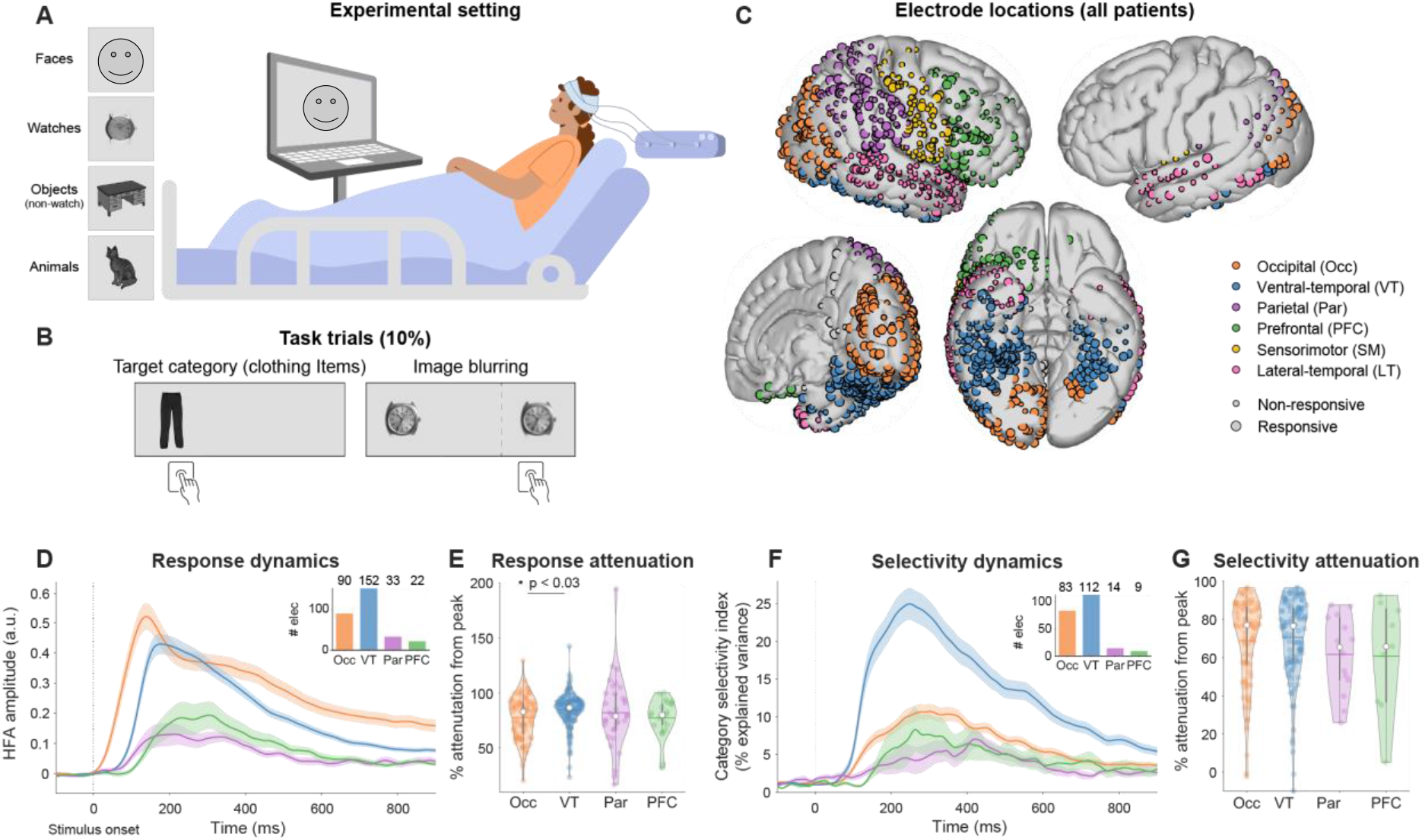
Experimental setup and design, electrode locations and single electrode response dynamics, showing substantial attenuation after the onset response. (A) Experimental setup and example images from the four categories. (B) Two target types (together 10% of trials): 1. Clothing items 2. Blurring of the image. (C) Electrode locations (pooled across patients), colored by ROI. The same color scheme applies for all figures. (D) HFA response dynamics relative to the pre-stimulus baseline (positively responding electrodes). To highlight response dynamics, for each electrode only trials from categories it was responsive to were included (inset: electrode numbers per region). Shaded area: SEM across electrodes. See also Figure S1A-E. (E) Relative attenuation in HFA responses from peak to 800-900ms (peak minus end activity relative to peak; attenuation >100% when end activity is lower than baseline levels). Colored dots: single electrodes, colored horizontal lines: means, white dots: medians across electrodes, gray vertical bars: interquartile range, contour lines: kernel probability density estimate. Black horizontal lines and asterisks: significant post-hoc differences between ROIs. (F) Category Selectivity dynamics in category selective electrodes (η^2^ expressed as percent explained variance from a one-way ANOVA between categories). See Figure S1F-H for electrode locations and single electrode properties. (G) Relative attenuation in selectivity, higher numbers indicate stronger attenuation. Notations as in (E). (D-G) image durations ≥ 900ms.

## Results

We measured broadband high frequency activity (HFA, 70-150Hz; Methods) shown to reliably track local neuronal activity^21,22^, in six a-priori regions of interest defined anatomically (ROIs; Figure 1C, Table S4). Of 907 noise-free electrodes, we focus on 430 which were visually responsive, defined as significant HFA modulation relative to a 200ms pre-stimulus baseline, for at least one category, in at least one of four non-overlapping 200ms time-windows between 100-900ms after stimulus onset (considering only images presented for 900ms or longer; see Methods and Table S5 for more details). Thus, electrode selection did not involve any category or temporal selectivity. Analyses focus on the four visually responsive regions: Occipital (Occ), Ventral-temporal (VT), Parietal (Par), and Prefrontal (PFC); other regions are shown in the Supplementary Information. With the exception of Figure 1D-E, all analyses in the main text include both positively (increased activity, >80% of sites) and negatively responding (decreased activity) electrodes. Similar results were obtained using positive electrodes only (Supplementary Information). To increase the number of trials in each category we pool all images presented for 900-1500ms for all analyses except for Figure 4 where we consider each stimulus duration separately.

### Response magnitude and category selectivity in single electrodes is substantially attenuated after the initial onset response

We first examined the single electrode HFA response dynamics in images presented for 900ms or longer (Figure 1D). Response magnitude was higher in Occ and VT relative to PFC and Par (one-way ANOVA, F(3,293) = 12.5, p < 10^-6^, Figure S1A), and peaked earlier in Occ relative to all other regions (F(3,293) = 15.73, p < 10^-8^, Figure S1B; see Figure S1C-E for response dynamics in other regions). Importantly, despite the continued presentation of the stimulus, response magnitude in all ROIs was substantially attenuated after the onset response, with ∼5-fold reduction in activity by 800-900ms after onset relative to peak response (mean attenuation ± SEM of electrodes from all four ROIs: 81.8% ± 1.1). Attenuation magnitude slightly varied between regions (one-way ANOVA, F(3,293) = 2.98, p < 0.033; Figure 1E) with post-hoc tests revealing only larger attenuation in VT compared to Occ (Tukey-Kramer method, p < 0.03, Cohen’s d = 0.42). VT and Occ showed strong responses at the onset, and even after the prominent response attenuation, their responses stayed (on average) higher than baseline levels (see our prior analysis of this data in ref. ^12^ for more detailed discussion of this point).

Of 430 responsive electrodes 236 showed significant category selectivity in at least one 200ms time-window (one-way ANOVA between categories; Methods, Figure S1F). In line with the general reduction in the magnitude of responses, by 800-900ms category selectivity (η^2^ values from temporally resolved ANOVA), declined by an average (± SEM) of 77.4 ± 1.1%, relative to peak selectivity (Figures 1F-G, S1G-H; no significant differences between regions, F(3,214) = 1.09, p > 0.3). Thus, both response amplitude and category selectivity in single electrodes were substantially attenuated after the onset response, despite the continued presence of the stimulus.

### Multivariate state-space dynamics in sensory regions track the duration of the stimulus

To understand how information is encoded in patterns of distributed activity, we examined the multivariate state-space trajectory in response to each category separately^23–25^ (Figures 2A, S2A; all images presented for 900ms or longer). We quantified multivariate activation using the point-by-point distance of the neural trajectory from the baseline state (when no stimulus was presented; insets). Similarly to the single electrode activation profiles, the multivariate activation increased rapidly after the onset, followed by marked attenuation (reduction of ∼80% from peak response to 800-900ms in all regions except Occ, Figure S2B). The decay of activity was substantially slower than the rise of the onset response, especially in occipital and ventral-temporal regions (see Figure S2C-D for analysis of state transition speeds). Separating the responses to stimuli of different durations (faces: Figures 2B, S3A; watches: Figure S3B) shows that despite the amplitude attenuation after the onset, multivariate responses in VT and Occ precisely tracked stimulus presence, with significant differences comparing 900 to 300ms, and 1500 to 900ms stimuli, emerging shortly after the offset of the shorter stimulus (permutation test with max-statistic control for multiple comparisons, p < 0.05). Responses in PFC and Par did not show this profile of duration dependence. Similar results obtained by comparing trajectories directly (without considering the distance to the pre-stimulus state; Figure S3C-D).

**Figure 2.**
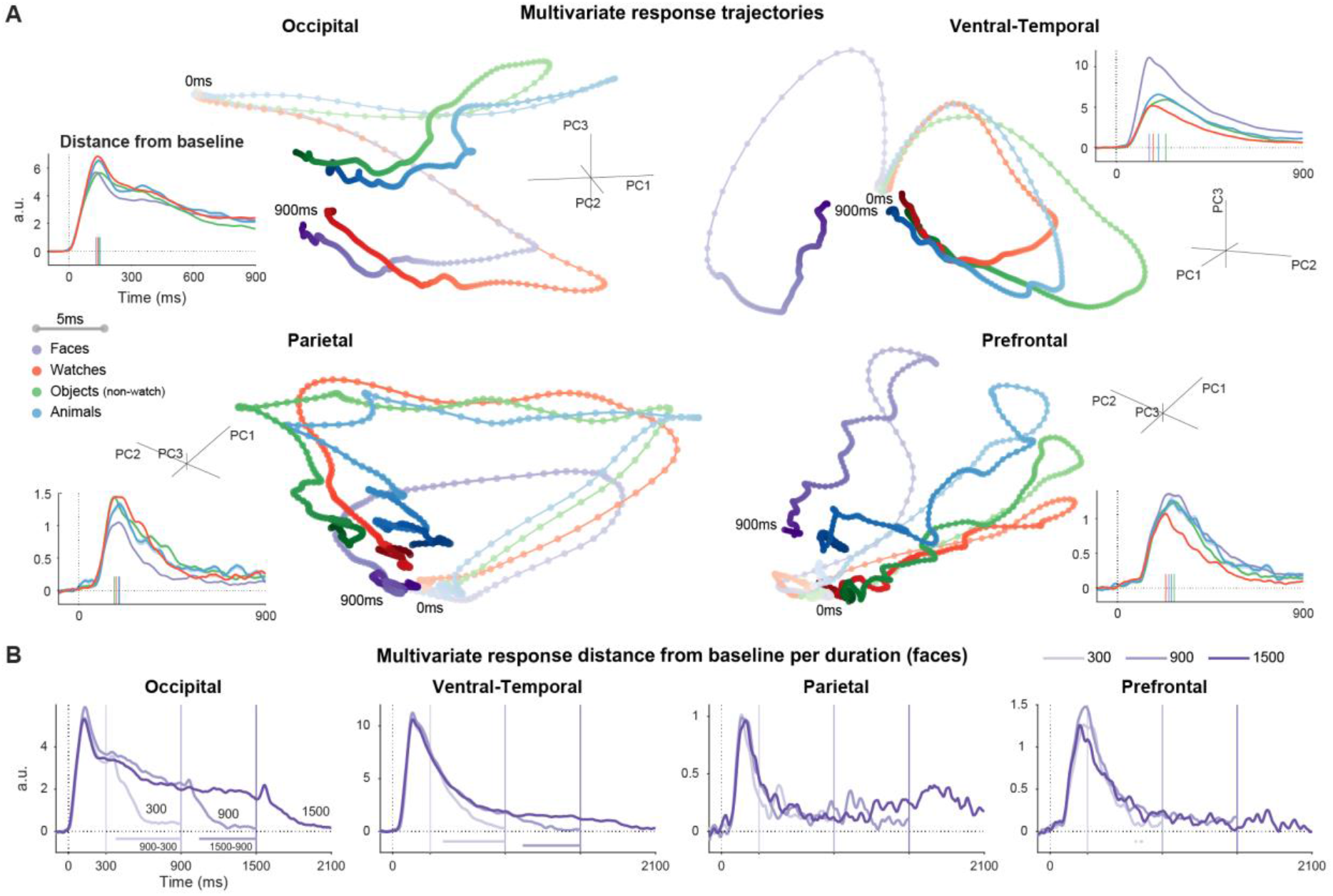
Multivariate state-space dynamics in sensory regions track the duration of the stimulus. (**A**) State-space trajectories per category (image durations ≥ 900ms; first 3 principal components using all responsive electrodes, responses in each category averaged prior to PCA; PCA was performed solely for visualization purposes). Trajectory lines are darker and thicker as time progresses, dots are 5ms apart. Insets: point-by-point distance of each trajectory from the pre-stimulus state (computed using the full response, prior to PCA); colored vertical lines on the abscissa: peak distance times. See Figures S2 and S3E for extended analysis of state-space trajectories. (**B**) Dynamics of distance from baseline (face images, see also Figure S3A; for other categories see Figure S3B; baseline subtracted for presentation purposes). Offsets marked by vertical lines with corresponding hues. Horizontal bars: time points of significant differences between durations (max-statistic permutations; 1500 vs. 900ms, 900 vs. 300ms; colors correspond to the shorter duration in the contrast). Traces are cropped 600ms after stimulus offset (shortest ISI). Absolute distances are comparable within region at different time-points, not between regions, as magnitude is dependent on the number of electrodes. See also Figure S3C-D.

### Visual category representation is sustained and stable in sensory regions and transient in frontoparietal regions

Next, we examine the representational content of the multivariate responses using time-resolved single-trial decoding^26,27^ on a subset of 92 unique images which were shown to all patients for 900ms or longer. Similar conclusions were obtained by examining the dispersion between state-space trajectories (Figure S3E). We trained linear classifiers for each ROI to distinguish between each pair of categories, using the HFA responses across electrodes as features (Figure 3A). Classifier performance was evaluated using area under the curve (AUC) of the receiver operating curve (ROC). Peak decoding was significantly higher than chance in both occipito-temporal and frontoparietal regions (Figure 3B; mean ± SEM across comparisons: VT, 99.6 ± 0.3%; Occ, 98.1 ± 1.6%; PFC, 84.1 ± 2.3%; Par, 86 ± 1.7%; one-sided max-statistic permutation test, VT and Occ all p_perm_ < 0.002, PFC and Par all p_perm_ < 0.041, except object-animal in both regions, and PFC_watch-object_). Thus, category information was not limited to traditional visual areas.

**Figure 3.**
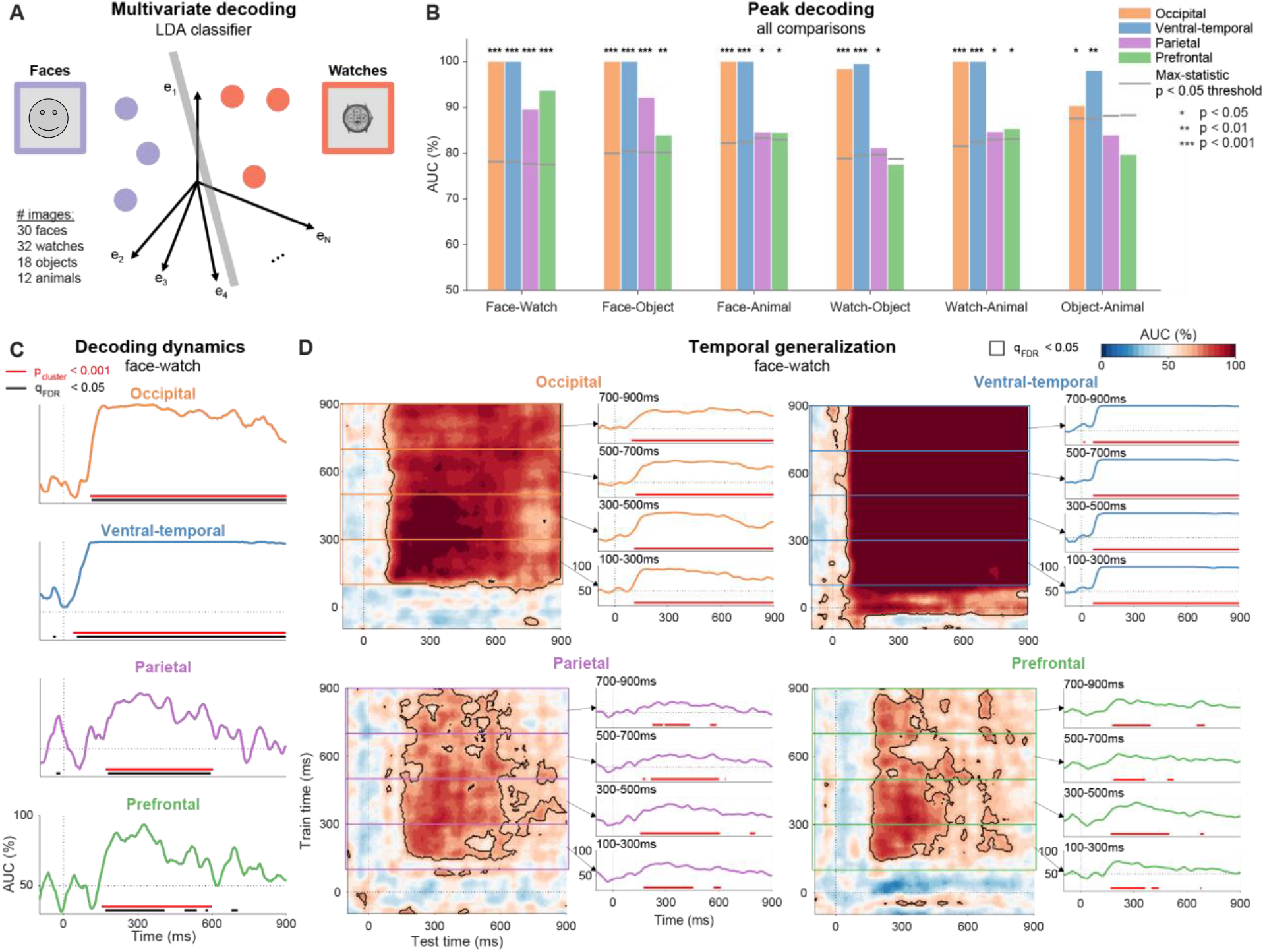
Visual category representation is sustained and stable in sensory regions and transient in frontoparietal regions. (**A**) Schematic illustration of decoding for a single time-point: colored dots represent single trial responses; a gray bar represents the linear classifier. (**B**) Peak decoding. Significance computed by permutation testing. Gray horizontal lines: significance threshold (max-statistic permutation testing; threshold is higher for comparisons involving categories with less exemplars). (**C**) Decoding dynamics (face-watch, other comparisons: Figure S4). Dashed lines: stimulus onset and chance level. Red bars: significant clusters by cluster-based permutations, black bars: significant points by point-by-point permutation testing (FDR corrected). (**D**) Temporal generalization matrices (face-watch, other comparisons are shown for VT and Occ in Figure S5D-E). The diagonal (training and testing on the same time-point) corresponds to the time-courses in (C). Black contour: contiguous points significant by point-by-point permutation testing (FDR corrected). Right-side plots: mean generalization dynamics for 200ms blocks of training time. Red bars: testing points significant for ≥ 50% of training points in the range. (B-D) All stimuli durations ≥ 900ms. See Figures S5-S6 for single patient results, analysis of Occ and PFC subregions, and control analyses ruling out the contribution of ocular muscle activity to PFC decoding.

**Figure 4.**
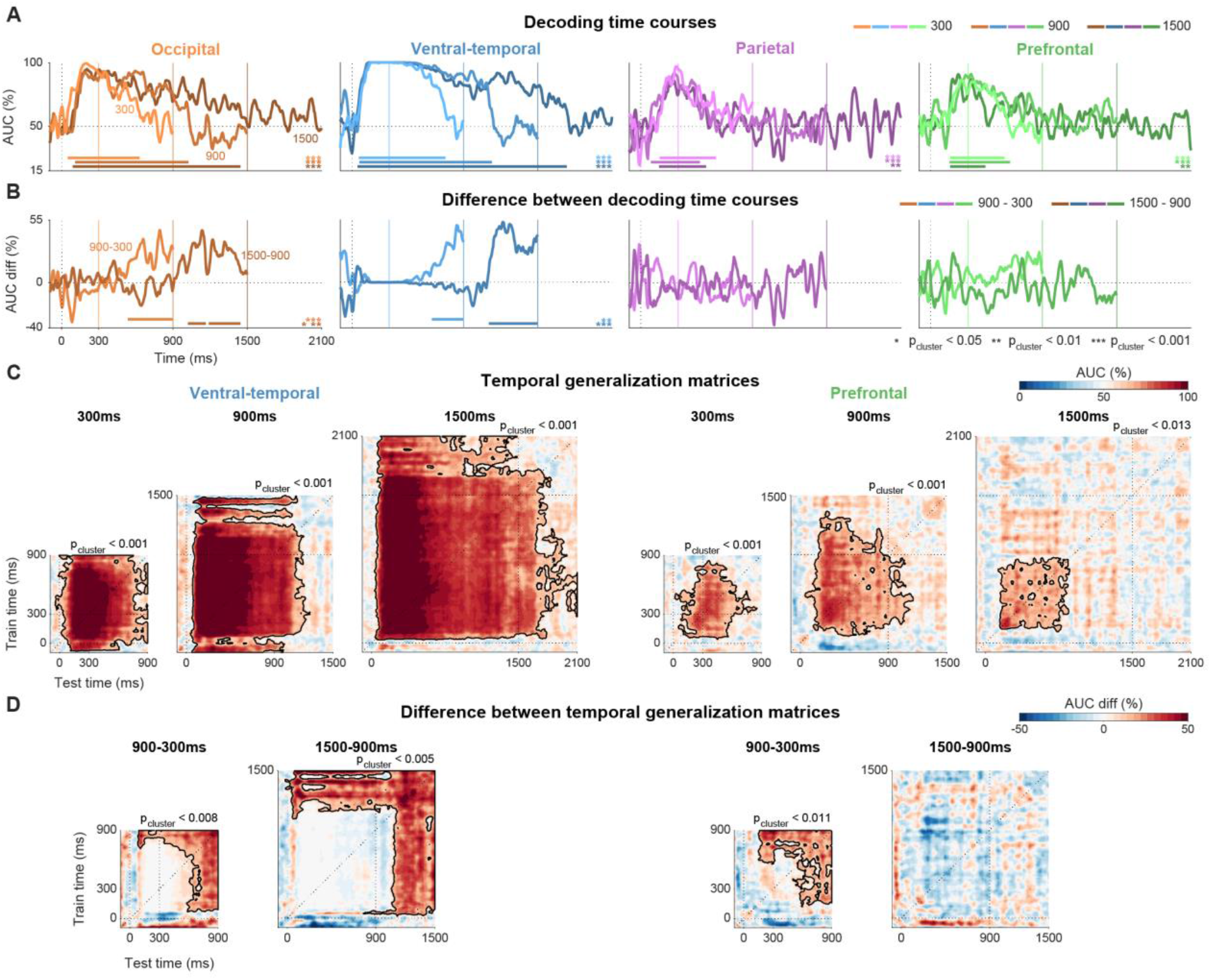
Category information in visual, but not frontoparietal regions, tracks stimulus duration. (**A**) Decoding dynamics per duration (darker lines correspond to longer stimuli, offsets marked by corresponding vertical lines). Horizontal bars of corresponding color: significant decoding clusters (cluster permutations); p-values are indicated by bottom-right corner asterisks (corresponding to the cluster temporal order). Traces are cropped 600ms after stimulus offset (shortest ISI). See also Figure S7A-E. (**B**) Difference of decoding time-courses (1500-900ms, dark lines; 900-300ms, bright lines). Statistical testing and notations as in (A). (**C**) TGMs per duration (see Figure S7F for the other regions). Dashed lines: stimulus onset and offset, and the diagonal (corresponding to the dynamics in (A)). Black contours: significant clusters; corresponding p-values shown above each TGM. (**D**) Comparison between durations (notation and statistical testing as in (C)). All panels: Occ and VT shown for patients S4-S10 (similar for S1-S3, Figure S7B). PFC is shown for S4-S10 as well (no responsive PFC electrodes for S1-S3). Par is shown for S1-S3 (not significant for S4-S10). See Methods and Tables S6-S7 for the rationale behind the split. All panels depict face-watch decoding (object-watch shown in Figure S7D).

However, the temporal profiles of decoding performance were dissimilar across regions (Figure 3C focuses on the face-watch comparison, all comparisons are shown in Figure S4A; direct time-resolved contrasts between regions are shown in Figure S4B). In VT and Occ, significant clusters^28^ emerged early after stimulus onset and persisted throughout stimulus presentation (all p_cluster_ < 0.001, red horizontal bars in Figure 3C; confirmed also with FDR corrected point-by-point permutations, black horizontal bars). Despite the substantial attenuation in response magnitude and in single electrode selectivity in both regions (Figure 1), category decodability decreased only minimally throughout this time (face-watch mean AUC 100-900ms: VT, 99.8%; Occ, 92.4%). These results did not stem from a single patient; high and persistent decoding performance was observed in the majority of patients with electrodes in these regions (Figure S5A-B). Similar results were found for both retinotopic and non-retinotopic regions of Occ (identified using a probabilistic map of visual topographic areas^29^; Figure S5C).

In contrast to the visual areas, significant category decoding in PFC and Par was transient, and mostly limited to ∼150-600ms after stimulus onset (face-watch, both p_cluster_ < 0.001; see Figure S6A-B for single patients). Onset times were delayed relative to the appearance of category information in sensory regions, consistent with the idea that content selective activity in frontoparietal regions only emerges after activity in sensory areas reaches a critical level^17^. We repeated the analysis separately for OFC and LPFC, as these parts of the cortex belong to partially distinct networks considering cytoarchitectonics, connectivity patterns and function^30,31^. Category information was significantly decodable from both subregions, though it was more prominent in OFC relative to LPFC (Figure S6C). Given the intense debate about the role of prefrontal representation in conscious awareness^17,32^, we applied several controls which ruled out the contribution of ocular muscle artifacts to PFC decoding (Methods and Figure S6D-E, as LPFC is less susceptible to ocular-muscle artifacts these analyses focus on OFC electrodes). Coverage in these regions was less comprehensive than in sensory regions (Figure 1C), which raises the possibility that the transient nature of category information in these regions stems from the reduced coverage. However, increasing the number of frontal electrodes by not limiting analysis to responsive electrodes led to similar results (Figure S4A). We also repeated the analysis in Occ and VT after reducing the number of electrodes to match coverage in PFC which nonetheless resulted in sustained category information in these regions (Figure S4C). Thus, PFC transiently represents category information even though no overt report was required for any of the stimuli used in the analysis^8,33^.

These findings show that visual areas provide reliable category information for as long as the image is presented, not only at the onset. This sustained decoding could stem from a series of changing yet discriminating patterns, or a single sustained state. To address this, we applied the temporal generalization method^34^, in which classifiers are trained on data from each time-point separately, yet each classifier is tested on *all* time-points, resulting in a temporal generalization matrix (TGM; face-watch: Figure 3D, other comparisons: Figure S5D-E). Successful decoding between time-points (off-diagonal decoding) indicates that the direction in state-space which discriminates between the categories remains stable in time. Thus, the rectangular temporal generalization pattern we reveal in occipito-temporal regions, and especially VT, indicates a highly stable, time-invariant category representation, as it shows classifiers trained during the onset response were able to distinguish between the categories during the sustained response and vice versa. This finding was also replicated at the single patient level (Figure S5A-B). In contrast to this temporal invariance in occipito-temporal areas, category information in PFC and Par did not generalize for the entire presentation of the stimulus (Figure 3D).

### Category information in visual, but not frontoparietal regions, tracks stimulus duration

The previous sections focused on responses to stimuli presented for 900ms or longer. To ensure the sustained decoding we found in visual sensory regions corresponds to the ongoing presence of the visual stimulus, and is not merely reflecting a prolonged onset response, we repeated the analysis separately for each duration (300, 900, 1500ms). To allow a large enough number of images for each duration and category, the analysis was performed separately on patients S1-S3 and S4-10 (see Methods and Tables S6-S7 for more details; Figure S7A shows electrode locations in each group). Category information in VT closely tracked the presence of the stimulus (with a short processing delay) – for all durations, significant AUC clusters emerged shortly after stimulus onset, persisted throughout stimulus presentation, then subsided with a delay of approximately 450ms for the 300ms stimuli, and 230ms for the two longer stimuli (face-watch comparison; Figures 4A and S7B-C). Occ showed a similar pattern, with somewhat shorter offset delays. In both regions, the direct time-resolved contrasts between decoding images in different durations (900-300, 1500-900ms) were statistically significant following the offset of the shorter stimulus (Figure 4B; all p_largest-cluster_ < 0.006). In contrast, decoding in Par and PFC did not correspond to the duration of the stimulus (time-course differences all p_cluster_ > 0.2). This conclusion is also supported by decoding of other category comparisons (Figure S7D) and by decoding using only positively responding electrodes (Figure S7E). This distinction between representational dynamics in sensory and frontoparietal regions was also evident by comparing the TGMs elicited by stimuli of different durations (VT and PFC, Figure 4C-D; Occ and Par, Figure S7F) – only occipito-temporal areas evinced time-invariant representations tracking the stimulus duration up to 1500ms, and this representation was stable in time.

To conclude, despite the drastic change in overall response amplitude (Figures 1-2), visual sensory areas maintain time-invariant category representations, tracking the duration of the stimulus. PFC and parietal cortex reveal reliable content representation, but only transiently and independent of stimulus duration.

### Exemplar representation is sustained and stable in sensory regions and transient in frontoparietal regions

Perception is more than recognizing categories. For example, when looking at a face, we do not merely see a face, we see a specific instance of a specific face. We therefore turned to examine exemplar-level information, using representational similarity analysis (RSA)^35,36^. RSA captures the representational structure by focusing on the pairwise dissimilarities between neural responses to each pair of images, grouped in a representational dissimilarity matrix (RDM). We quantify neural dissimilarities as 1-Pearson correlation, but with few exceptions we note, similar results were obtained using Euclidean distance dissimilarity (Figures S8-S9). We consider a region as representing exemplar information if the representational structure, conveyed by the dissimilarities, is reliable across separate repetitions of the same exact stimuli. Thus, we focus on 60 images viewed by five of the patients at least twice (for 900ms or longer, as in previous analyses). Results from a larger group of eight patients (with only 18 shared images) are shown in Figures S8-S9 (see Methods for more details and Figure S8A for electrode locations in each subgroup).

To examine the reliability across repetitions we designed two complementary metrics, both computed on a time-point by time-point basis. Item Reliability (IR) captures the reliability of each stimulus representation within the overall geometry of responses, by measuring separately the reliability of the response to each stimulus relative to other stimuli (Figure 5A). Geometry Reliability (GR) captures global aspects of the representational structure by comparing the full dissimilarity structure across repetitions (Figure 5B; averaging different pairs of Geometries or correlating each pair separately before averaging led to similar results). Note that both IR and GR compare dissimilarities between and within repetitions. Thus, by design, they capture not only preservation of the dissimilarity structure, but also of state-space location (see Methods for more details). Finally, both metrics are tested against surrogate distributions generated by shuffling stimulus identity (z-scoring shown in Figure 5A-B) and thus both metrics also reflect discriminability of single exemplars by the geometry (if multiple exemplars occupy similar positions within the geometry shuffling them will not alter the geometry, and this will reduce the IR and GR scores).

**Figure 5.**
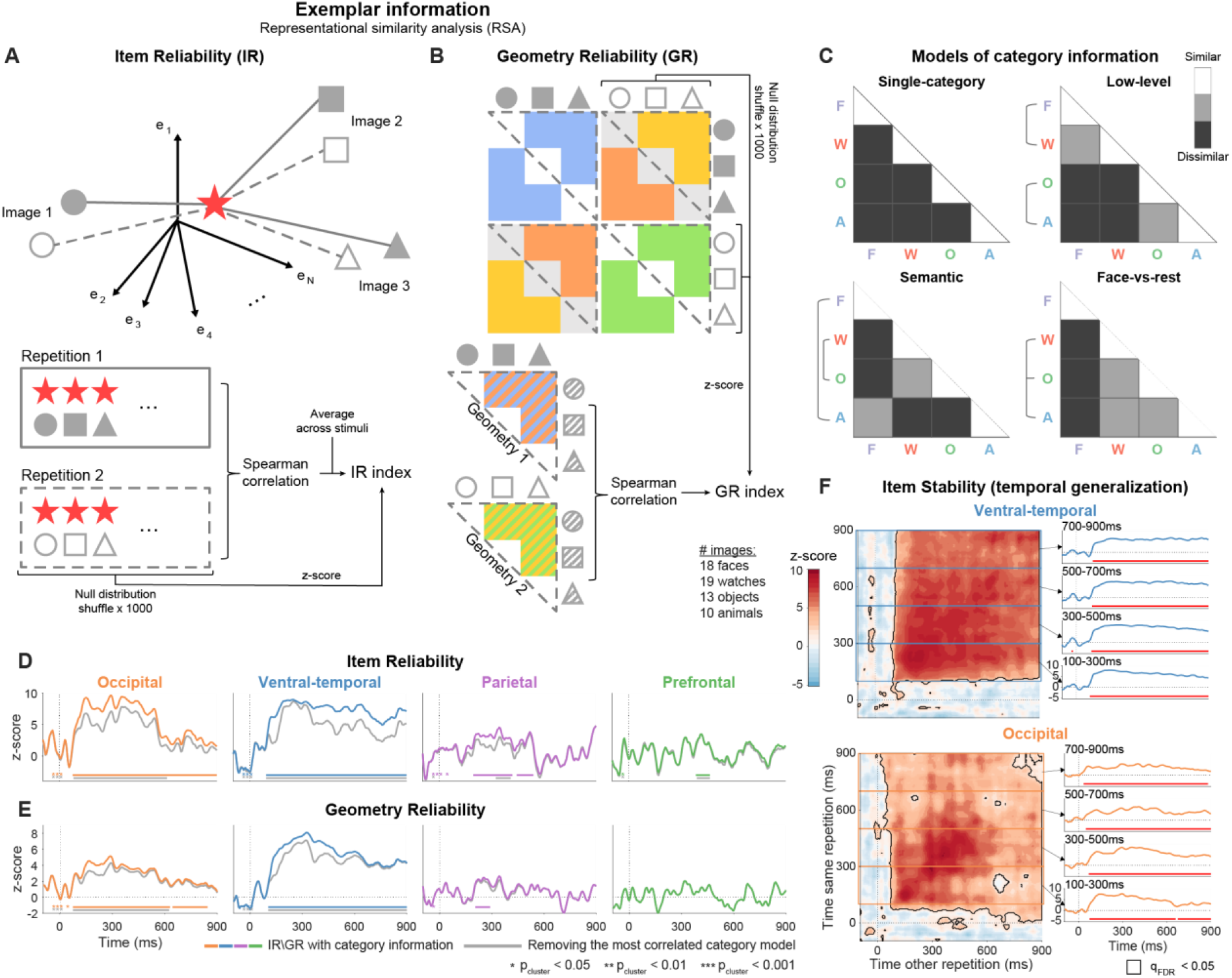
Exemplar information is sustained and stable in sensory regions and transient in frontoparietal regions. (**A**) Schematic illustration of Item Reliability (IR): for each image (red star) we compare the vector of dissimilarities to all other images in Repetition 1 (full shapes) to the vector of dissimilarities to all other images in Repetition 2 (empty shapes). (**B**) Geometry Reliability (GR): we first compute the dissimilarities between all images in both repetitions, resulting in a symmetric matrix with four distinct representational structures (top). Pairs of geometries are averaged to yield Geometries 1 and 2, and the two geometries are correlated. See Methods for more details about both reliability metrics. (**C**) Models of potential category information in the representational geometry. All models assume exemplars within each category are similar to each other and dissimilar to other categories. Three of the models add a hierarchy of similarity between categories (Methods). (**D**-**E**) IR and GR dynamics. Colored lines: full representational geometry; gray lines: after partialling-out the model explaining the most category information (see Figure S10 for more details about the calculation and removal of other category models; Figure S8 for other control analyses). Horizontal bars of the same color mark significant clusters; p-values indicated by bottom-left corner asterisks (corresponding to the cluster temporal order). Dashed lines: stimulus onset and chance level (no single-item information). (**F**) IR temporal stability (GR: Figure S9B, removing category information: Figure S10D-E; other controls: Figure S9C-E). Notations as in Figure 3D. (D-F) Images presented at least twice with duration ≥ 900ms.

Starting with VT, we found highly reliable exemplar representation, sustained throughout stimulus presentation (IR and GR clusters extending to 900ms, p_cluster_ < 0.001; Figure 5D-E colored lines, see Figure S8B-C for additional regions not shown in the figure). We further tested for temporal invariance of the representation by comparing dissimilarities between time-points (analogous to decoding temporal generalization) and found highly stable representation (high off-diagonal stability; IR: Figures 5F, GR: Figure S9B; see S9A for additional regions). To verify that our reliability metrics reflect coding of single exemplars, rather than relying solely on the category structure that we observed previously (Figures 3-4), we repeated the calculation of both metrics after removing category structure from the representational geometry. This was done by partialling-out from the neural RDM four models of potential category information, designed in accordance with the literature and the observed state-space trajectories (Figure 5C, see Methods for more details). The gray lines in Figure 5D-E depict IR and GR dynamics after partialling-out the model which was most strongly correlated with the RDM of the region (Figure S10A), thereby providing the strictest measure of category information (using other models led to similar or higher reliability scores, Figure S10; see also Figure S11 for exemplar reliability within single categories supporting the same conclusions). Both IR and GR remained sustained and stable in VT throughout stimulus presentation after controlling for category information (Figure 5D-E gray lines, Figure S10B-E). Using only positively responding electrodes, using Euclidean dissimilarities, or using data from the eight-patient group led to similar results (Figures S8D-F and S9C-E).

In Occ, exemplar representation was largely sustained and stable using correlation dissimilarity (both p_cluster_ < 0.001), but this result was less robust: representation was not fully sustained nor stable after removing the categorical structure, nor with Euclidean dissimilarity or the larger patient group (Figures 5D-E, S8D-F and S9C-E). Exemplars were also reliably represented in frontoparietal cortex, though this effect was short-lived and noisier than in visual sensory regions (IR significant in both regions, all p_cluster_ < 0.015; GR significant only in Par, p_cluster_ < 0.043, albeit not after accounting for category structure). Thus, we conclude that representation of exemplars is sustained and stable (time-invariant) in visual sensory regions, though it is robust only in VT, and there is transient and weaker exemplar representation in PFC and Par.

## Discussion

Delineating the neural correlates of conscious experience is one of the most coveted yet most challenging goals of cognitive neuroscience and perhaps science at large^1^, leading to multiple competing hypotheses^16,37^. In different guises, the quest for the NCC typically involves looking for an isomorphism between a specific experience and neural signal by contrasting two states: having an experience (being consciously aware) of a stimulus and not having one^33,38^. Usually, this requires unnatural manipulations of the stimuli, for example by masking, or by major manipulations of attention^39^. Under these liminal conditions, observers sometimes experience stimuli and sometimes not, and neural signals are then compared. These are powerful paradigms, especially for examining processing without conscious awareness, but in many cases determining whether a stimulus was genuinely not experienced is difficult and when experience is present, it is typically impoverished due to the manipulation^40^. As our results show, temporal manipulation of presentation duration of visible stimuli provides an illuminating alternative, as neural correlates of experience should correspond to the full temporal extent of experience, not only the experience of onsets or changes which previous studies focus on.

By examining which aspect of the neural response corresponds to the duration of stable perceptual experience, we addressed a glaring gap in the quest for the NCC, and vision neuroscience in general – how do we maintain a stable experience beyond the immediate onset of a stimulus, when neural activity, as well as local selectivity, drops dramatically^12,13^? We show that despite variability in response magnitude (Figures 1-2), the pattern of activation across recording sites in occipital and especially ventrotemporal cortex contains sustained information about the visual percept at both the category (Figure 3) and exemplar level (Figure 5), and this representation is stable (invariant) in time, lasting for the entire the duration of the stimulus (Figure 4). These properties are commensurate with the introspective intuition of ongoing, continuous perceptual experience. In contrast, we found a burst (‘ignition’) of visual information in PFC and parietal cortex, even though no report was required. This representation lasted for a few hundred milliseconds after onset and did not correspond to the duration of the stimulus. This suggests that frontoparietal regions may be involved in updating perceptual experience, including when no overt response is needed, and is commensurate with a more discrete aspect of perceptual experience^4,5^.

### The temporal structure of experience

Introspectively, consciousness feels like a continuous flow of experiences, progressing in “real-time” with events in the environment. However, this prevalent intuition has been challenged on both empirical and philosophical grounds^3,41^. First, neural transmission takes time, and moreover, processing delays vary between modalities, and even between different features in the same modality, complicating the relation between perception and the external world. Second, perception of changes, motion, and melodies, require integration of a temporal interval, rather than a momentary instance. This conclusion is also supported by the existence of postdictive effects, when a presented stimulus alters the way we perceive *prior* stimuli^2,42^. One well known example is the “Color Phi phenomenon”: two differently colored discs are flashed sequentially in different positions on the screen, but despite the discrete nature of the stimuli, the experience is of a disk moving between the two locations and changing color midway. Both the direction of movement and the color are fully unpredictable before the appearance of the second disk, thus, the perception of movement during the time interval between the two flashes must be a retrospective “filling in” mechanism. One prominent proposal to resolve this puzzle is postulating discrete perception^4,5^ – if no perception at all occurred before the second flash, the full perceptual event can be organized unconsciously and experienced without any inconsistencies. However, the mechanism behind these types of effects is still studied, and continuous solutions have also been proposed^3^.

In the context of our paradigm, the introspective subjective experience is of *stable* continuous percepts with varying durations. Taken at face value, this would suggest that the sustained and stable visual representations underpin our ongoing conscious experience. However, to the extent that perception is composed of discrete samples, each generating a transient ignition, the frontoparietal representation would correspond more directly to experience. Thus, distinct representational dynamics, mapping to distinct hypotheses about the temporal nature of experience, seem to coexist in different regions. Rather than being mutually exclusive, these representations may be related to different aspects of a multifaceted experience. The sustained and the onset representations may interact hierarchically to form our ongoing conscious experience (see ref. ^42^ for one such proposal). Taken together, the results show a deep connection between the classical NCC problem, which largely emphasized the anatomical underpinnings of consciousness, and the longstanding debate about the temporal nature of awareness.

It should be noted that our stimuli durations (up to 1.5 sec) were substantially longer than the gaps between perceptual samples typically posited by theories of discrete perception (typically ∼0.1 sec, or at the extreme end, 0.5 sec)^4,5^, which would predict finding additional bursts of content information in PFC beyond the onset. Yet, if these repeated ignitions are not temporally aligned between trials (except for an onset ignition) they may be missed by our analysis, resulting in a time-course with only a transient onset representation. Finally, our results do not rule out a continuous role of PFC in monitoring or supporting this representation in a manner that the does not represent the specific content of visual experience^32^.

### The role of prefrontal cortex in perceptual awareness

Prefrontal involvement in perceptual awareness was criticized recently by several studies which reported minimal or no PFC activation when subjects are not required to respond overtly or report awareness of the stimuli^43,44^ (so called “no-report paradigms”^8^). These findings suggest that PFC is not involved in experience per se, but in reporting it (though see ref. ^45^ for a recent no-report study which did find PFC awareness effects). Unlike the aforementioned no-report paradigms, the trials we analyzed, while not requiring a response, were task relevant, as the subjects had to decide whether the stimulus belonged to a target category. Nevertheless, our findings are unique in showing human stimulus specific representations which are not associated with an overt report, and are not mapped to task distinctions^46^. That is, we find highly accurate decoding between image categories that are non-targets in the PFC. Moreover, the task only required discrimination of category information, yet we find also reliable exemplar representation. Thus, contrary to recent claims that prefrontal cortex manifests only non-specific task-related activity^47^, our results support content representation in PFC following the onset of a new stimulus (for related findings in monkeys see ref. ^48,49^).

### Implications for specific theories of consciousness

An ongoing adversarial collaboration (COGITATE), aiming to adjudicate between two prominent theories of consciousness, IIT and GNWT, has adopted our multi-duration paradigm as one of two key tests^16,50^. An important contribution of this project has been the delineation of detailed predictions by the two theories regarding this paradigm. IIT predicts that neural activity in the “posterior hot-zone”, including occipital and VT regions, will persist with a stable representation as long as the visual experience persists, while it is largely agnostic about prefrontal involvement. GNWT predicts an ignition (transient onset representation without a sustained component) in PFC starting 250-300ms after stimulus onset, including when no overt behavioral responses are performed. Both predictions in fact bore out in our study – posterior (visual) areas showed stable persistent representation, tracking the duration of the stimulus, and PFC showed an ignition without persistence, absent behavioral responses. Thus, these predictions of IIT and GNWT are, in fact, not adversarial. Rather, the persistent representation in occipitotemporal cortex and transient representation in frontoparietal cortex may tap onto different components of experience, or to different levels in a hierarchical view of time-consciousness^42^.

The parietal cortex has been traditionally linked with the prefrontal cortex by theories of conscious awareness, based on findings that changes in perceptual content, like in binocular rivalry^51^, or detection of liminal, masked stimuli (relative to no detection)^52^, activate both parietal and frontal regions. Thus, GNWT posits that parietal cortex is included in the “global workspace” alongside PFC^17^, and accordingly, predicts transient representation in the region. More recently, proponents of IIT delineated the concept of the “posterior hot-zone”, predicted to manifest sustained and stable temporal dynamics. The exact anatomical boundaries were not precisely stipulated but limited diagrams and verbal descriptions of the hot-zone typically include parietal cortex alongside occipitotemporal regions^47,53,54^. We find that parietal cortex displays transient, onset-related visual representations, which do not persist for the duration of the stimulus (Figures 4A-B and S7F). Thus, parietal responses resemble the PFC responses much more than the occipital and ventrotemporal cortex responses, in line with the prediction of GNWT. That said, the parietal lobe is far from being a homogenous structure^55^, and different parts of the parietal lobe may manifest different response dynamics.

### Distributed representation and the “Experience Subspace”

The notion that perceptual representations in humans are distributed, rather than local, has been suggested mainly based on fMRI findings^56,57^, and remains contested^58^. The temporal stability of distributed representation reported here, in face of substantial local variability, provides strong support to the importance of multivariate patterns. This finding is consistent with the “neural population doctrine”, which posits that the fundamental unit for neural computation and cognition is a distributed ensemble rather than a single neuron^23–25^. A shift from identifying the NCC with activation in specific neurons, to the multivariate population response, is also supported by a recent binocular rivalry study in monkeys showing that the same neurons in high-order visual cortex which represent the perceived stimulus also code the suppressed image simultaneously. Yet, a decoder trained on the population response was able to closely track the ongoing percept, indicating that even in the face of local heterogeneity, conscious content can be reliably coded at the population-level^59^.

The representations we identified in ventral-temporal cortex were not only sustained, but also stable. That is, the same classifier could discriminate categories from the onset response to the end of the presentation time in occipitotemporal cortex, and similarly, the same representational geometry persisted throughout this time. Our results thus reveal an “Experience Subspace” within the vast space of possible neural responses, which maintains consistently the distinctions between visual categories and between exemplars within each category, despite considerable moment-by-moment variability in the state-space response. We propose that to the extent representations in sensory regions contribute to our conscious experience, it is the projection of the population response to this subspace which imbues each perceptual experience with its unique quality, while the variable activity in other dimensions may be used for other functions or may simply be the result of neural stochasticity. The stability within the “Experience Subspace” also affords downstream regions with a stable readout of information about the visual percept^60^. This notion of stable subspace also helps avoid the need to invoke multiple realizability^61^, a perennial conundrum in the philosophy of mind regarding whether a specific conscious experience can be realized by more than one state (see also ref. ^62^). Our analysis goes one step further, as in addition to stable stimulus information we also identified reliable time-invariant coding of the relational structure between neural responses to different exemplars (Figure 5). This relational stability is consistent with the recent proposal that the subjective quality of an experience is derived from the relation of its neural representation to other previously learned representations^63,64^. This may also explain the perplexing dissociation between our perception of repeating stimuli as identical, despite the ubiquitous phenomenon of repetition suppression^65^ – we predict that as long as the response to a stimulus occupies the same location within the subspace, it will be similarly perceived regardless of the initial or sustained amplitude of response.

A related proposal has been put forward to explain the puzzling phenomenon of “representational drift”, that is, the observation that tuning properties of single neurons vary dramatically between trials or sessions of testing, separated by seconds to many days, even in the face of consistent behavioral performance^66,67^. This dissociation between stable behavior alongside representational drift resembles our findings of perceptual stability in the face of dramatic attenuation of the neural response, albeit at a different temporal resolution. Most of the studies of representational drift involved single unit recordings or local ensembles of neurons in laboratory animals, and our findings extend the notion to humans and more broadly distributed representations (see also a recent fMRI finding of representational drift across sessions^68^). In the context of representational drift, several authors have suggested that drift is confined to a coding “Null Space”, that is, it influences population activity orthogonally to coding of task-related variables and will therefore not interfere with task performance^67,69–71^ (but see ref. ^72,73^). Importantly, studies of representational drift typically treat representation as stable at the sub-second scale, yet our results show considerable response variability within a single event, therefore more work is needed to connect the two phenomena.

## Conclusion

Which parts of the brain reflect our current perceptual experience and how their temporal dynamics correspond to the subjective experience are two major questions in the quest for understanding the neural correlates of conscious awareness. By manipulating stimulus duration, we were able to identify a hitherto undescribed duality between these two aspects. In sensory regions, we find sustained and stable representation in an “Experience Subspace”, embedded within the variable, diminishing neuronal responses. In frontoparietal regions we found discrete (transient) content representation at stimulus onset. Thus, if the introspective subjective experience of a continuous stable percept given stationary input is accurate, it is the invariant sensory representation within the “Experience Subspace” which imbues each perceptual experience with its unique, consistent quality. Yet, to the extent consciousness is discrete, this could be aligned with the discrete transient prefrontal representation. Thus, understanding the temporal resolution of experience will shed light on the anatomical location of the NCC, and the NCC will in turn inform our understanding of the temporality of conscious experience.

### Limitations of the study

First, the prefrontal cortex is a heterogeneous structure, with complex spatial organization^30,31^, and mixed selectivity at the single neuron level^74^. Persistent information could be present in sites not sampled by our electrodes which were placed solely based on clinical considerations, and in this study concentrated mostly on one hemisphere. The lack of sustained representation could also be related to the relatively small number of electrodes placed in the PFC, though reducing the number of electrodes in sensory regions to the same number available in PFC did not substantially deteriorate the sustained representation observed in those regions (Figure S4C). PFC may still be involved in maintaining sustained continuous percepts using activity in low frequency bands, or in silent synaptic changes^75^. For all these reasons, the absence of (sustained) activity should be taken with more caution than the presence of activity (as is always the case with null results). Second, the stimuli analyzed in this study were all above threshold, clearly visible images. Thus, it is possible that unseen subthreshold images are similarly encoded over time. Future studies should test this possibility by comparing representation of seen and unseen sustained images, and directly manipulate the duration of aware visual experience independent of the duration of the input, as in binocular rivalry^59^. Finally, our use of the term “representation” is meant to denote neural patterns which correlate with external stimulus descriptors and should not be understood as implying mechanistic or functional roles^76^. Nor do we claim that the representation used by the brain is constrained by our ROIs, which were chosen to adhere to common divisions of the brain, and to address arguments from the different consciousness theories.

## Acknowledgments

We are grateful to members of the Human Cognitive Neuroscience Laboratory (HCNL), and A. N. Landau and the Landau lab for insight and support throughout the study, R. Malach and R. Broday-Dvir for helpful discussions, and H. T. Vishne for multiple late nights of assistance with figures. We thank patients who participated in the experiment and J. Parvizi, R. A. Kuperman, K. I. Auguste, P. Weber, K. Laxer, D. King-Stephens and E. F. Chang for enabling data collection from their patients. This work is dedicated to the memory of Mrs. Lily Safra, a great supporter of brain research. G.V. is supported by the Azrieli foundation graduate fellowship. L.Y.D. is supported by the Jack H. Skirball research fund. This study was supported by US-Israel Binational Science Foundation grant 2013070 (awarded to L.Y.D. and R.T.K.) and National Institute of Neurological Disorders and Stroke grant 2 R01 NS021135 (R.T.K.).

## Author contributions

G.V. and L.Y.D. conceptualized and designed the study and methodology. L.Y.D and E.M.G. designed the experiment, L.Y.D., E.M.G. and R.T.K. collected and curated the data. G.V. performed formal analysis, model development, programming, and visualization. G.V. and L.Y.D. wrote the manuscript, R.T.K. reviewed it. L.Y.D. and R.T.K. provided funding for the study. L.Y.D. supervised the study.

## Declaration of interests

L.Y.D. is the co-founder and shareholder of, and receives compensation for consultation from Innereye Ltd., a startup neurotech company. The company business is not related to the current study. The authors declare no competing interests.

## STAR★METHODS

### KEY RESOURCES TABLE

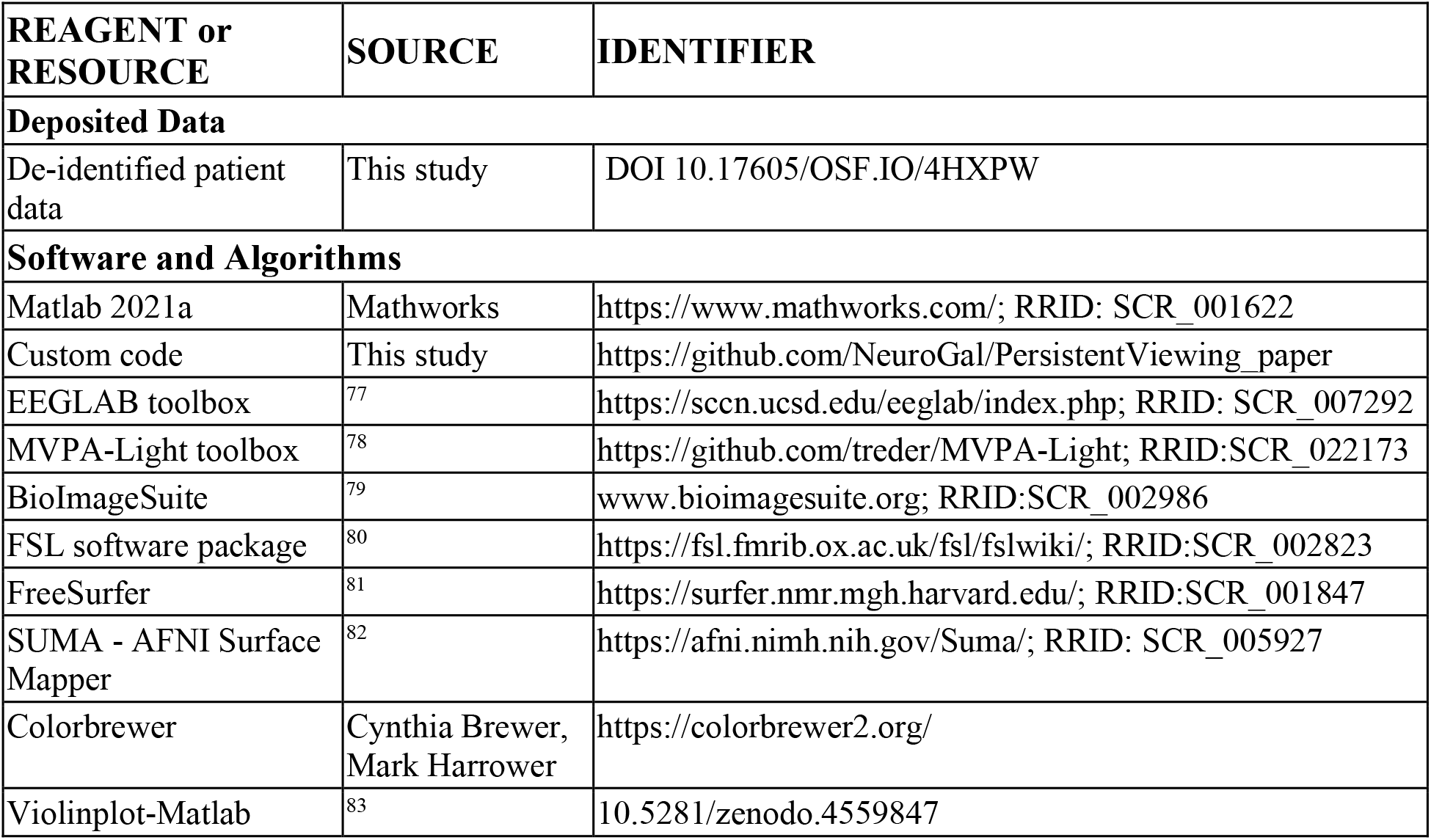

### RESOURCES AVAILABILITY

#### Lead contact

Further information and requests for resources should be directed to and will be fulfilled by the lead contact, Gal Vishne.

#### Materials availability

The study did not generate new unique reagents.

#### Data and code availability

- De-identified human data have been deposited at OSF. They are publicly available as of the date of publication. DOIs are listed in the key resources table.
- All original code used for analyses and visualizations has been deposited at Github and is publicly available as of the date of publication. DOIs are listed in the key resources table.
- Any additional information required to reanalyze the data reported in this paper is available from the lead contact upon request.

### EXPERIMENTAL MODEL AND SUBJECT DETAILS

Ten patients undergoing presurgical evaluation for treatment of intractable epilepsy (4 female, age (mean ± SEM): 41 ± 3.7, range: 19-65; for individual demographic details see Table S1). Recordings were conducted in the Epilepsy Monitoring Unit (EMU). Seven patients were recorded in the Stanford School of Medicine, two in the California Pacific Medical Center (CPMC) and one in the University of California, San Francisco Medical Center. All patients gave informed consent approved by the University of California, Berkeley Committee on Human Research and corresponding IRBs at the clinical recording sites, in accordance with the Declaration of Helsinki. Results from the same dataset were previously reported in ref. ^12^.

### METHODS DETAILS

#### Stimuli and task

Patients viewed grayscale images, presented at the center of a uniform gray background, and extending approximately 5° of the visual field in each direction. Stimuli were presented on a laptop screen and responses captured on the laptop keyboard (Figure 1A). The images belonged to multiple semantic categories, including faces (∼30%), man-made objects (watches: ∼30%, other objects: ∼18%) and animals (∼10%). ∼10% of images were targets (see below). The remaining images (<3%) were mostly houses or body-parts, which were not analyzed due to paucity of exemplars (see Table S2 for the number of stimuli viewed by each patient). Watches and other man-made objects were considered separately in this study since watch images, like the face images, were taken from a dataset of photos, while the other objects were illustrations. When comparing to face images, watches are considered a better control over low-level similarities, as the two categories share the overall round outline with internal details^20^. To verify that information content tracked ongoing stimulus presentation, images were presented for variable durations (patients S1-S3: 300, 600, 900, 1200 or 1500ms, S4-S10: 300, 900 or 1500ms). The probability of each exemplar appearing in each duration was uniformly distributed. A fixation cross was displayed between image presentations (inter-stimulus interval: 600, 750, 900, 1050 or 1200ms). Image sequence was randomized for each patient, meaning that specific exemplars were viewed for different durations by different patients.

Patients were instructed to fixate at the center of the screen and respond with a button press to the appearance of rare targets (Figure 1B). In the main experimental condition, performed by all patients, the target was any image of a clothing item. The second experimental condition was performed by seven patients (S4-S10) in half of the blocks. In this condition patients were instructed to respond to two target types: (1) appearance of a clothing item (as in the main condition), (2) blurring of any image during the last 200ms of presentation (see ^12^ for more details on the motivation of the dual task). Both target types together comprised ∼10% of the trials and were not analyzed in this study, focusing on trials without report. Neural responses to non-targets did not differ between conditions^12^, therefore data from both conditions was analyzed together.

#### Data acquisition and preprocessing

Patients were implanted with 64-128 subdural electrodes (total 1004), arranged in 1-dimensional strips and\or 2-dimensional grids (AdTech Medical Instrument Corporation). Electrodes were 2.3 mm in diameter, with 5 or 10 mm spacing between electrodes. Eight patients were implanted in the right hemisphere, and two in the left (Table S3 for individual electrode coverage). Two patients were additionally implanted with depth electrodes (total 28 electrodes), which were not analyzed in this study. We excluded an additional 35 channels which did not record any signal. Recordings were sampled at 1000Hz (CPMC), 3051.76Hz (Stanford, UCSF) or 1535.88Hz (Stanford) and resampled to 1000Hz offline. A high-pass filter was applied online to the signal at either 0.1Hz (five patients, increased to 0.3Hz for parts of the recording in two of the patients), 0.16Hz (one patient) or 0.5Hz (four patients). Electrodes manifesting ictal spikes or persistent noise were identified visually and removed from further analysis (0-38 electrodes per patient, total 125; only analyzed electrodes are plotted in visualizations of electrode positions). Electrodes were re-referenced offline to the average potential of all noise-free electrodes (per patient). Line noise (60Hz and harmonics) was removed offline by a custom made notch filter, designed to remove persistent oscillations (not transients)^84^. All data processing and analysis was done in Matlab (Mathworks, Natick, MA) using custom code or the toolboxes referenced in the key resources table. Visualization was done using custom code except violin plots which were created using ref.^83^. Colormaps were created using colorbrewer (https://colorbrewer2.org/) with some adaptations.

#### Electrode localization

Electrodes were localized manually using BioImageSuite^79^ on a post-operative Computed Tomography (CT) scan co-registered to a pre-operative MR scan using the FSL software package^80^. Individual patient brain images were skull-stripped and segmented using FreeSurfer^81^. Localization errors (resulting from co-registration errors or anatomical mismatch between pre-and post-operative images) were reduced using a custom procedure which jointly minimizes the squared distance between all electrodes within a single electrode array or strip and the cortical pial surface. Individual patients’ brains and electrode coordinates were co-registered to a common brain template (FreeSurfer’s fsaverage template) using surface-based registration^85^, which preserves the mapping of electrode locations to anatomical landmarks, and each cortical surface was resampled to a standardized mesh using SUMA^82^ (see ^12^ for more details). Cortical electrodes were assigned to one of six anatomical regions of interest (ROIs) based on the FreeSurfer automatic parcellation (ventral-temporal, occipital, prefrontal, parietal, sensorimotor and lateral-temporal; Figure 1C and Table S4). Prefrontal electrodes were further divided into lateral-prefrontal and orbitofrontal cortices, and occipital electrodes were divided into retinotopic and non-retinotopic based on a probabilistic map of visual topographic regions^29^ (see electrode locations in Figure S1D). Twelve noise-free electrodes located over medial regions (mostly in the precuneus or cingulate cortex) were excluded from further analysis due to their paucity. Visualization of electrode positions was based on surface registration to an MNI152 standard-space T1-weighted average structural template image.

#### High-frequency activity estimation

We focus analysis on high-frequency activity (HFA, 70-150Hz), previously shown to track firing rate in humans^21,22,86^ and other primates^87,88^. We excluded the low-gamma range used in a previous study with this data, as it was shown to manifest distinct spectral and functional properties^89,90^. To estimate the HFA time-course we band-pass filtered the whole signal in eight 10Hz sub-ranges between 70-150Hz (EEGLAB’s FIR Hamming window, function ‘pop_eegfiltnew’^77^). We then extracted the instantaneous amplitude in each band using the Hilbert transform and normalized by dividing the signal by the mean amplitude in that range. Finally, we averaged the amplitude traces from all bands. Normalization was done to account for the 1/f profile of the power spectrum, which results in reduced contribution of the high frequencies relative to the lower frequencies. Trial segments were defined from -300ms to 1600ms around each stimulus onset, and baseline corrected by subtracting the mean HFA signal in the 300ms prior to stimulus onset from the entire trial segment. Trials containing excessive noise were excluded from analysis. The resulting HFA time-courses were smoothed by a 50ms moving window (smoothed time-courses are used in all analyses unless noted otherwise).

### QUANTIFICATION AND STATISTICAL ANALYSIS

In the following sections we describe the four parts of the analysis in detail: Single electrode responses, Multivariate state-space responses, Decoding of category information, and Exemplar specific information. Following the description of the dependent measures in each section, we describe the approach to statistical analysis. Note that we use a multiverse approach, testing hypotheses in multiple ways to ensure the robustness of the results^91^.

#### Single electrode responses

##### Visual responsiveness

Responsiveness was tested separately for each of the four categories (considering watches and other objects separately), in four non-overlapping “stimulus-on” windows: 100-300ms, 300-500ms, 500-700ms and 700-900ms after stimulus onset, considering only trials with durations of 900ms or longer. For each category and each time-window, we compared the mean HFA signal during the “stimulus-on” window to the mean HFA signal 200ms prior to stimulus onset (*two-tailed* paired t-test; averaging was done prior to smoothing the HFA trace to avoid information leakage between windows). We used Bonferroni correction across windows, and FDR correction (Benjamini-Hochberg procedure^92^) across electrodes, thus, electrodes with q_FDR_ < 0.05/4 in at least one of the four windows were considered responsive to that category. As the test was two-tailed, electrodes were considered responsive both when activity during the “stimulus-on” window was increased relative to the pre-stimulus window, and when it decreased. Changes in response sign between windows or categories were rare (<5% of responsive electrodes), thus, electrodes were classified as positively responding (increase from baseline; 346/430 electrodes; Figures 1D-E, S1A-D) or negatively responding (decrease from baseline; Figure S1E) based on the sign of the sum of t-statistics from all tests (across all categories and all time-windows). Both types of electrodes were used in all analyses except where noted.

##### Response latencies and post onset attenuation

We first computed the mean HFA response of each electrode (using only stimuli from categories which the electrode was responsive to with stimuli duration of 900ms or longer). Peak response magnitude (Figure S1A) and peak response time (Figure S1B) were defined as the maximal HFA value between 0-900ms after onset and the time-point when this was achieved, respectively.

Relative attenuation of response magnitude was defined as the difference between the peak response and the mean response 800-900ms after onset, scaled by the peak response (Figure 1E):

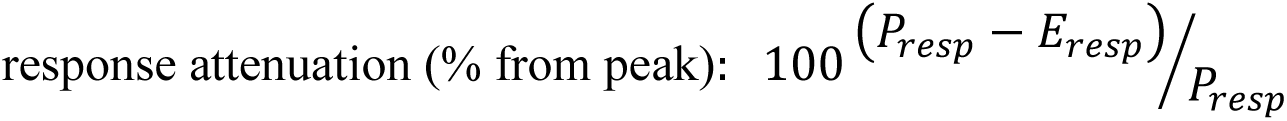

where P_resp_ is the peak response magnitude and E_resp_ is the mean response 800-900ms after onset.

##### Category selectivity

Electrodes were defined as category selective if they showed differential responses to stimuli of the four categories in at least one of the four windows used above for stimuli longer than 900ms (one-way ANOVA, Bonferroni corrected across time-windows and FDR corrected across electrodes, similarly to the responsiveness criterion; Figure S1F). For each category selective electrode, we defined a selectivity time-course by extracting the percent of variance in responses explained by category information on a point-by-point basis (100 ⋅ 𝜂^2^, where 𝜂^2^is the effect size measure from a point-by-point one-way ANOVA; Figure 1F-G, S1G-H).

#### Multivariate state-space responses

The main multivariate analyses (including decoding and exemplar analyses) were performed grouping together responses from multiple patients (c.f. ^93,94^). Yet, to ensure no result is driven by a single patient we repeated the central analyses in single patients and found highly consistent results (Figures S5-S6).

##### Analysis of state-space trajectories

The multivariate neural state at a specific time instance 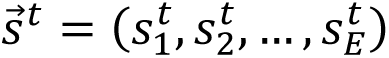 is defined as the pattern of recorded activity across electrodes^23–25^ (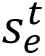 is the response of electrode e at time t and E is the number of responsive electrodes in a region). State-space trajectories record the changing multivariate states across time. To quantify the multivariate response magnitude (distance from baseline; Figures 2, S2A and S3A-B) we computed the point-by-point Euclidean distance of the trajectory from neural state prior to stimulus onset. Since all trials were baseline corrected, this is equivalent to computing the L^2^ norm of the vector of responses across electrodes. Attenuation was calculated similarly to the univariate case (Figure S2B).

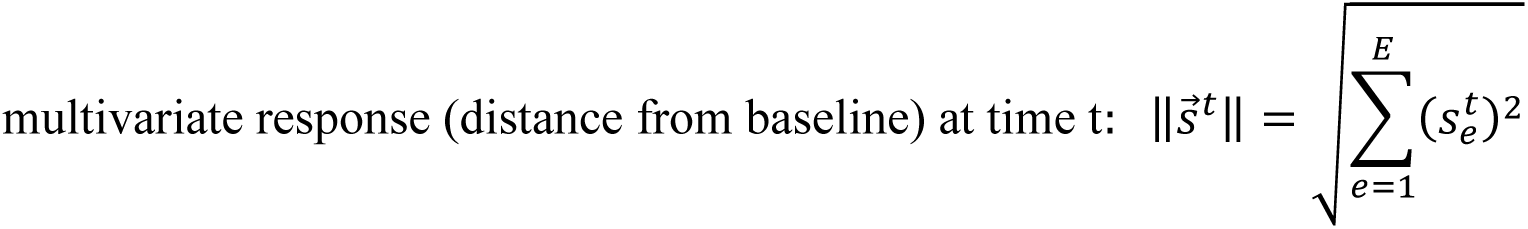

State transition speed (Figure S2C-D) was quantified as the distance travelled by the neural state-space trajectory in 1ms (corresponding to our sampling rate).

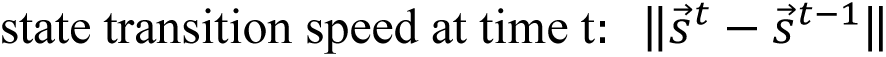

To quantify whether the neural trajectories tracked the duration of the stimulus we computed the time-point by time-point distance between the responses to stimuli of different durations (Figure S3C-D).

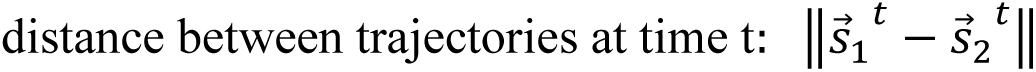

where 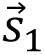 and 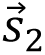 are response trajectories to different stimulus durations.

To evaluate category information in the multivariate trajectories (Figure S3E) we calculated the dispersion between response trajectories to different categories: First, we computed the square of the Euclidean distance (L^2^ norm squared) of each trajectory from the mean response across all categories. We defined the category selectivity index as the square root of the mean of these distances (analogous to computation of standard deviation in a one-dimensional distribution), and z-scored relative to a permutation null distribution as described in the next section.

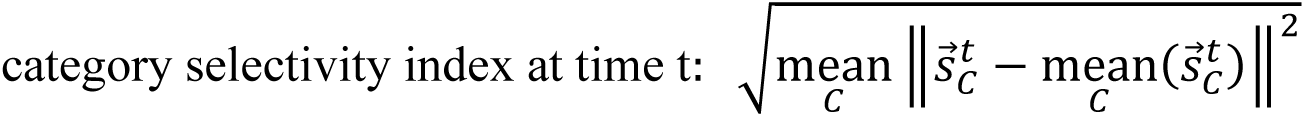

where 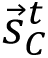 is the neural response to category C at time t, and 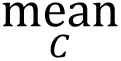 is the average over all four categories.

##### Statistical testing and confidence intervals

To evaluate statistically the dependence of multivariate response trajectories on stimulus duration (Figures 2B and S3A-D) we used permutation testing with max-statistic control for multiple comparisons^95^ (N_perm_=1000, separate permutations for 900-300 and 1500-900 comparisons). Since stimulus duration is expected to influence the response only at the times when one stimulus is still presented and the other is not, we considered only time-points between 300 and 900ms for the former comparison and between 900 and 1500ms for the latter (results were highly similar considering the entire time-course). In each permutation we shuffled the duration labels across trials, before averaging the trials of each duration to construct the surrogate duration-specific state-space trajectory and computing the relevant difference statistic for each time-point. Thus, we created N_perm_ permutation statistic time-series (N_perm_ x N_time_ matrix). Next, we computed the mean and standard deviation across permutations for each time-point (column means and standard deviations) and used these to standardize (z-score) all permutation time-series. To create the surrogate distribution, we extracted from each standardized permutation time-series the maximal z-value across all time-points. We also z-scored the values of the unpermuted (true label) statistic in the same manner, using the means and standard deviations of the permutation matrix (as under the null hypothesis the true statistic comes from the same distribution as the permutation statistics). Finally, time-points when the unpermuted z-scored statistic was larger than 95% of the null distribution were considered significant (one-sided test). The procedure to test for significant category information (Figure S3E) was similar, except that we shuffled the category affiliation across trials and all time-points were considered in the calculation.

Confidence intervals (Figure S2B,D) were computed using a jackknifing approach^96^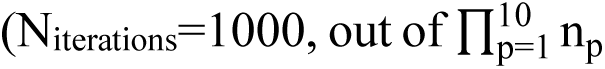 possible combinations per category, where n_p_ is the number of trials of that category viewed by patient p). In each iteration, for each patient, the state-space response trajectory to each category was estimated by averaging the response to n_p_ - 1 trials (separately for each region). We then merged the response trajectories of single patients (each with its own electrodes) to form a single response trajectory with all electrodes in each cortical ROI. Percent attenuation and speed difference for each iteration were computed in the same way as for the neural response calculated based on all images, and the distribution across iterations is provided as the confidence interval.

#### Decoding category information

##### Time-point by time-point classification

To quantify the representation of category information in each cortical region, we trained for each pair of categories, a set linear classifiers (one per time-point) to distinguish between trials when one category was presented (e.g., faces) vs the other category (e.g., watches), using the HFA amplitude of all responsive electrodes in the region as features^27^ (Figure 3A). For all analyses under this section (Figures 3-4 and S4-S7) the data was downsampled to 200Hz to reduce computation time. To avoid overfitting, decoding analyses were carried out using five-fold cross validation, and to minimize variability stemming from the stochasticity in fold assignment, we repeated the five-fold cross validation procedure five times and averaged the results. In each iteration of the calculation, the category containing more trials was undersampled to balance the number of trials in both categories and we ensured that both categories were represented roughly equally in all folds.

We used regularized Linear Discriminant Analysis (LDA)^97^, as implemented in the MVPA-Light toolbox^78^ (downloaded on 04/02/2021). LDA attempts to find a one-dimensional projection of the data (a linear weighting of electrodes) with maximal separation between the categories. This is done by simultaneously trying to maximize the distance between the mean responses to each category (“signal”), while minimizing the variability in responses to each of the categories (“noise”). Formally, classifier weights (the projection vector) are computed according to:

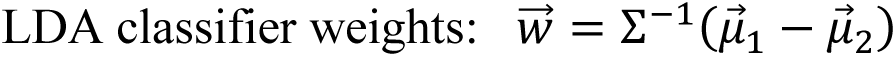

where 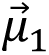 and 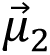 are the mean responses to category 1 and category 2 (e.g., faces and watches), and Σ is the pooled covariance matrix (summing covariances of each category, weighted according to the number of exemplars per category). The predicted category for each trial at each time-point is given by comparing 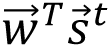 to a set threshold θ:

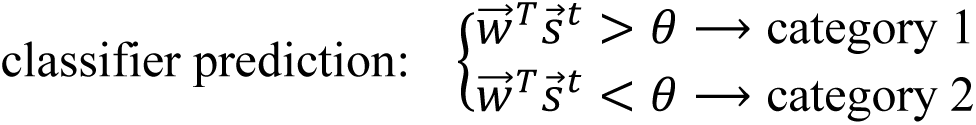

The threshold can be modified to alter the balance of true positives (trials from category 1 classified as category 1) and false positives (trials from category 2 wrongly classified as category 1), resulting in the receiver operating curve (ROC), which depicts the false positive rate (FPR) and the true positive rate (TPR) for each threshold θ. We quantified the classifier’s performance by using the area under the ROC curve (AUC):

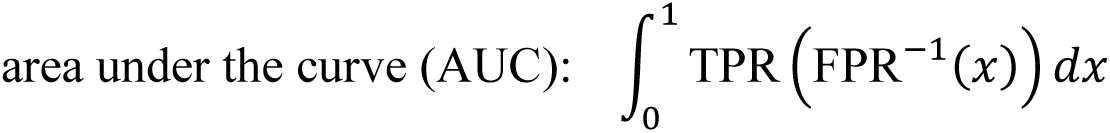

When the classifier contains no category information the AUC is equal to 0.5 (50%, chance level performance), and when the classes are fully separable the AUC is equal to 1 (100%, perfect performance). We used AUC, as opposed to accuracy, since AUC is more robust to unequal class sizes, and is not influenced by changes in the overall magnitude of the response when the pattern across electrodes remains similar.

To reduce the influence of random noise fluctuations on the weight estimate we used a shrinkage estimator for the covariance matrix. Instead of using empirical covariance in the equation above, we use a combination of the empirical covariance (Σ) and the identity matrix (I), weighted according to the regularization parameter λ:

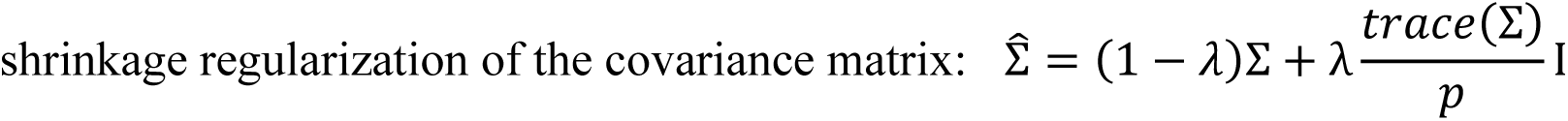

where p is the number of features of the classifier (equal to the number of responsive electrodes). Multiplying the identity matrix by the mean value on the diagonal ensures that the trace of the covariance is preserved, which helps to mitigate the bias introduced by this step. The regularization parameter λ was estimated using the Ledoit-Wolf formula^98^ (default implementation in MVPA-Light^78^, for more explanation on the rationale of shrinkage regularization see ^99^).

The main decoding analyses were carried out on responses to 92 unique images which were seen by all patients for ≥ 900ms (faces, 30; watches, 32; other objects 18; animals 12; Figures 3B-D, S4, S5C-E and S6C). To avoid overfitting^27^, repetitions of the same image were averaged to ensure it is not used simultaneously in the training and testing sets (number of repetitions mean ± std across patients and unique exemplars: 1.5 ± 0.6, range 1-4, similar for all categories). Using only the first repetition of each image instead of averaging the repetitions elicited highly similar results. Analyses performed on data from single patients used all unique exemplars viewed by each patient for ≥ 900ms (mean ± SEM unique exemplars across patients: faces, 62.5 ± 4.2; watches, 65.5 ±

4.5; objects (non-watch) 39 ± 2.3; animals, 22 ± 1.1; Figure S5A-B, S6A-B,E). Comparison of bipolar and average montages was performed on patients S8 and S10 together (faces, 54; watches, 58; other objects, 34; animals, 20; Figure S6D). To run analyses for each stimulus duration separately (Figures 4 and S7) we split the patients into two groups (S1-S3, S4-S10), in order to increase the number of exemplars seen in the same duration by all patients in the group (due to the randomization the full group shared on average only 4.8 exemplars per category and duration). The number of exemplars in each category and duration shared by the full patient group and each sub-group are shown in Table S6.

##### Control for potential ocular muscle artifacts

Previous studies have shown that HFA activity in the proximity of the orbit may be confounded with saccade related activity from extraocular muscles (spike potentials)^100,101^. While these studies explicitly showed no contamination of responses in OFC (as opposed to the temporal pole)^100^, and patients were instructed to fixate during our task, we nevertheless performed two control analyses to ensure that the ability to decode category information from OFC was not driven by eye-muscle activity.

###### Decoding using bipolar reference montages

(Figure S6D): We re-referenced the data after extraction of HFA traces by subtracting from each electrode the mean HFA activity in all adjacent electrodes on the same grid. Both responsive and non-responsive electrodes were included for re-referencing but only contacts centered on responsive electrodes were used for classification. Two patients had enough electrodes placed over OFC (inset above Figure S6D): S8: 16 OFC electrodes, 5 excluded (4 placed on arrays which were mostly over the temporal pole; 1 containing epileptic activity), resulting in 11 electrodes (5 responsive); S10: 10 OFC electrodes (6 responsive) + 2 LPFC electrodes on the same grid (used for re-referencing).

###### Decoding in time-windows without any saccades

(Figure S6E): Eye-tracking was not possible in the EMU setting, yet we were able to reliably identify the timing of saccades in one patient (S8), by detecting the saccadic spike potentials in an electrode placed over the temporal pole, behind the right eye (same electrode used in ^12^). Following ref. ^84^, we first convolved a template for the saccadic spike potential with the signal from the relevant electrode (for the template we used the validated publicly available “matched filter”^84^). Second, we marked all points higher than 3 standard deviations of the entire convolution time-course. Finally, we denoted the onset of each marked time-range as a saccade. This produced the expected saccade modulation curve around stimulus onset (suppression followed by rebound; similar to Figure 8 in ^12^), supporting the assertion that the detected time-points correspond to saccadic events. It was not possible to exclude all trials where a spike potential was detected between 0 and 900ms, as this would exclude 90% of the relevant trials and leave just 1-2 unique images in the animal and (non-watch) object categories. To overcome this problem, we focused on four non-overlapping time-windows between 100ms and 900ms after stimulus onset and analyzed each window separately while excluding only trials where a saccade occurred within that time-window. Using this approach, we were able to use 67-82% of the images across time-windows (72-78% of images from each category). Decoding analysis was carried out on the mean activity in each time-window (unsmoothed data), resulting in one AUC value for each time-window and category comparison. We used only responsive OFC electrodes which were not adjacent to temporal pole electrodes.

##### Temporal generalization analysis

To assess whether coding of category information was stable in time we used the temporal generalization method^34^ (Figures 3D, 4C-D, S5 and S7F). In this method linear classifiers are trained on data from each time-point separately, but tested on data from all time-points, not only the one used for training. The result is a matrix of decoding values (Temporal Generalization Matrix, TGM), with the y-axis indicating the training time point and the x-axis indicating the testing time-point. Successful decoding between time-points indicates that the direction in state-space which discriminates between the categories remains stable in time (the specific cutoff between categories may change as we used AUC to evaluate classifier success, which does not rely on one fixed threshold).

##### Statistical testing

All statistical testing was based on permutation tests (N_perm_=1000; one-sided). To construct the null permutation distribution, in each permutation we permuted the category labels of all trials and trained classifiers for all time-points based on the permuted labels (we used the same set of permuted labels to preserve the temporal structure of the data). Peak decoding performance (Figure 3B) was statistically tested by comparing to the distribution of peak AUCs of each permutation (maximum across time-points), equivalent to a max-statistic approach to control for multiple comparisons^95^. Controlling for multiple comparisons for decoding time-courses and TGMs was done using cluster-based permutations^28^ (details below; Figures 3C, 4, S4A-B, S5A-C, S6A-D and S7B,D-F). Cluster-based permutations are sensitive, yet they are insufficient to establish the precise latency or temporal extent of effects^102^, therefore, we additionally employed point-by-point comparisons controlling the FDR^92^ whenever inferences about specific time-points were required (q_FDR_ < 0.05; Figures 3C-D, S5 and S7C).

Cluster-permutation details: We selected all time-points with AUC greater or equal to 60% (first-level threshold; other thresholds led to similar results), clustered the samples based on temporal adjacency (including both train and test times for TGMs) and extracted the sum of AUC values in each cluster (cluster statistic). To construct the null distribution, we performed the same procedure on each of the permutations, but only the maximal cluster value was retained in the null distribution. Clusters from the original (non-permuted) calculation that exceeded 95% of this distribution were considered significant (one-sided test).

Statistical comparison of decoding between regions (Figure S4B) and comparison of decoding of stimuli in different durations (Figures 4 and S7) was performed in a similar way, except the cluster first-level threshold was set to AUC difference of 10% between duration conditions. For comparison between regions we used a two-sided test. In both cases we permuted the category labels and trained classifiers for each region (or duration) separately and formed the null distribution by subtracting the decoding results for different regions, or, in the duration case by subtracting the short duration (300 or 900ms) from the long duration (900 or 1500ms) and extracted the cluster statistics as described above. When we compared the decoding time-courses between durations we considered only time-points after the offset of the short stimulus, as we predicted a difference only when one stimulus was presented, and the other was not (results were nearly identical without this constraint). For TGMs we considered the full matrix as we were also interested in generalization between the onset and the time-period after the offset of the short stimulus.

#### Exemplar specific information

##### Quantifying exemplar representation

To quantify exemplar-level information we used representational similarity analysis (RSA)^35,36^, which describes neural representation in terms of the relation between neural responses, that is, the geometry that the responses define. The representational geometry is fully captured by the set of all pairwise dissimilarities between the pattern of responses across electrodes to each pair of stimuli, grouped together in a representational dissimilarity matrix (RDM). We employed two dissimilarity measures – correlation dissimilarity (Figures 5, S8B-D,F, S9A-C,E, S10-S11) and Euclidean dissimilarity (Figures S8E, S9D), as these are sensitive to different aspects of the response^103^:

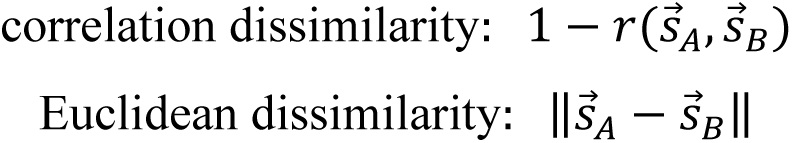

where 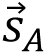 and 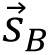 are the vector of HFA responses to stimulus A and B across electrodes (in a specific time-point), 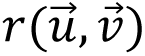 is the pearson correlation of 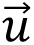 and 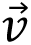 and ‖⋅‖ is L_2_ norm.

To avoid overfitting to incidental noise fluctuations, exemplar-level analyses were only conducted on stimuli which were viewed at least twice (akin to a cross-validation procedure), for 900ms or longer. Four patients viewed only 27 exemplars twice from the analyzed categories, and one patient did not view any exemplar twice, thus, to ensure sufficient sampling of all categories we performed the main analyses in this section on a group of five patients who all viewed the same set of 60 unique images at least twice (faces, 18; watches, 19; other objects, 13; animals, 10; Figures 5, S8B-E, S9A-D, S10-S11). Results from a larger eight patient group (excluding the patient who did not view any exemplars twice and one patient which had excessive noise in many trials) with only 18 unique exemplars (faces, 6; watches, 7; other objects, 2; animals, 3) are presented as a control in Figures S8F and S9E. Electrode locations of both groups are shown in Figure S8A.

We computed the dissimilarity structure separately for each repetition and time-point and compared between repetitions (reliability analysis) and between time-points (stability analysis) using Spearman correlation, which is more robust to changes in the overall magnitude of activity relative to Pearson correlation. As with the decoding analysis, for all analyses under this section (Figures 5 and S8-S11) the data was downsampled to 200Hz to reduce computation time.

##### Representation reliability across repetitions

###### Item Reliability

(IR; Figures 5D,F, S8B-F, S9A,C-E, S10B,D, S11): If exemplars are represented reliably across repetitions, the location of each exemplar within the neural geometry should remain consistent across repetitions. Thus, for each stimulus presentation (Figure 5A, red star), we computed the correlation of the vector of dissimilarities to all other images in repetition 1 (filled shapes) and the vector of dissimilarities to all other images of repetition 2 (empty shapes). We then averaged the obtained correlations across all images and repetitions, and z-scored the result using a permutation null distribution (N_perm_=1000). To construct the null distribution, for each image presentation we shuffled the stimulus identity of all images from the *other* repetition and repeated the process (i.e., if the presented image was from repetition 1 (as in Figure 5A) we shuffled repetition 2 and if the presented image was from repetition 2, we shuffled repetition 1).

###### Geometry Reliability

(GR; Figures 5E, S8B-F, S9, S10C,E): Our second measure of reliability captures global aspects of the representation by comparing the full dissimilarity structure across repetitions. We first computed the RDM between all stimulus presentations from both repetitions (Figure 5B, top), containing four distinct sets of dissimilarities: within repetition 1 (blue), within repetition 2 (green) and two between repetition 1 and repetition 2 (yellow and red). Second, we paired each within repetition dissimilarity set with one between repetitions dissimilarity set and averaged the dissimilarities in each pair (resulting in Geometry 1 and Geometry 2 in Figure 5B bottom; averaging different pairs of geometries or correlating each pair separately and then averaging led to similar results). Considering both within repetition and between repetition dissimilarities ensures not only preservation of the geometry across repetitions, but also a similar state-space location between repetitions. Third, we computed the correlation between Geometry 1 and Geometry 2 (unfolded into vectors). Finally, we z-scored the result using a permutation null distribution (N_perm_=1000), constructed by shuffling the identity of all *single exemplars* in repetition

2 and repeating the procedure. Importantly, in cases where some exemplars are represented similarly within the geometry (indistinguishable representations) this approach is likely to result in insignificant GR even if the geometry is fully maintained between repetitions. Thus, it is a test of both representational reliability between repetitions and of discriminability between exemplars.

##### Representation stability in time

We tested the temporal stability of the representation by comparing representational structures across time-points (Figures 5F, S9, S10D-E, S11C). This was done similarly to the reliability analyses, only representational structures were computed in different time-points. IR stability for time points (t_1_, t_2_): for each stimulus presentation S, we first computed the dissimilarities between S at time t_1_ and all other exemplars in the *same* repetition as S also at t_1_ (y-axis of stability plots), and then correlated this dissimilarity vector to the dissimilarities between S at time t_2_ and all other exemplars in the *other* repetition at t_2_ (x-axis of stability plots). We then averaged the correlation across stimuli as in the standard IR calculation. GR stability for time points (t_1_, t_2_): we correlated Geometry 1 in t_1_ (y-axis) with Geometry 2 in t_2_ (x-axis). When t_1_ = t_2_ (diagonal of stability matrices) this is equivalent to calculation of IR and GR, respectively. Note that by correlating dissimilarity structures at different repetitions (for at least one of the exemplars), we avoid overfitting to spontaneous fluctuations unrelated to stimulus processing, which are known to exhibit many temporal dependencies^104–106^.

##### Accounting for category structure

To test whether exemplar-level information is present beyond category differences we designed four models of category information (Figure 5C) and tested whether exemplar-level information is reliable and stable after removing category information from the neural RDM using partial correlation (i.e., partialling-out the model RDMs; Figures 5D-E, S10). For IR we used a single row of the model RDM for each stimulus (treating the RDM as a symmetric matrix, but excluding the diagonal), and for GR we partialled-out the full model RDM unfolded into a vector. All models assume exemplars within a category are similar to each other, and dissimilar to exemplars in other categories. In models (2-4) we add a hierarchy of similarities between categories (supported by previous studies^57,107,108^), such that categories belonging to the same higher order category are 50% similar. The models are (Figure 5C): (1) “Single-category” – no relation between categories, only a primary category distinction. (2) “Low-level” relation between categories based on low-level visual similarity – faces and watches form one high-order category (both are photos of round objects), and other (non-watch) objects and animals form the second (pointy illustrations). (3) “Semantic” (based on animacy) – faces and animals form one high-order category (animate) and all objects (watches and non-watches) form the second (inanimate). (4) “Face-vs-rest” – faces are distinct from all other categories, which are all grouped into one high-order category; we constructed this model due to the unique social importance of faces and the known specialization in face representation (also supported by our results, Figure 2A).

##### Statistical testing

All statistical testing was permutation-based (N_perm_=1000). Construction of the null distribution is detailed above (section ‘Representation reliability across repetitions’, different procedures for IR and GR). In each permutation, we used the same set of permuted labels for all time-points to preserve temporal properties of the data. Z-scoring was performed in each time-point separately, for both the original (non-permuted) score and the permutation results. Statistical testing and control for multiple comparisons was done similarly to the decoding analyses (all one-sided): Time-courses were tested using cluster-based permutations^28^ (sum of reliability indices as the cluster-statistic). First-level threshold was set to z = 1.5 SD for the main analyses (Figures 5D-E, S8, S10B-C) and z = 0.75 SD for the single category analyses (adjusted to accommodate the larger temporal smoothing window in these analyses (100ms vs 50ms in all other analyses); Figure S11B-D), in both cases similar results were obtained with other thresholds. The time-courses of correlation between each category RDM model and the neural RDM (Figure S10A) were also corrected using cluster-based permutations, with the sum of the correlations as the cluster-statistic and ρ=0.0391 as the first-level threshold (corresponding to a p-value of 0.05). Stability (generalization) matrices were tested using point-by-point comparisons, controlling FDR^92^ (q_FDR_ < 0.05; Figures 5F, S9, S10D-E).

## Notes

### Competing Interest Statement

LYD is the co-founder and share holder of, and receives compensation for consultation from Innereye Ltd., a startup neurotech company. The company business is not related to the current study.

### Summary of Updates

Broad updates in both the main text and the supplementary

