## Supplementary information for "Representing experience over time: sustained sensory patterns and transient frontroparietal patterns"

### *Supplementary Figures*

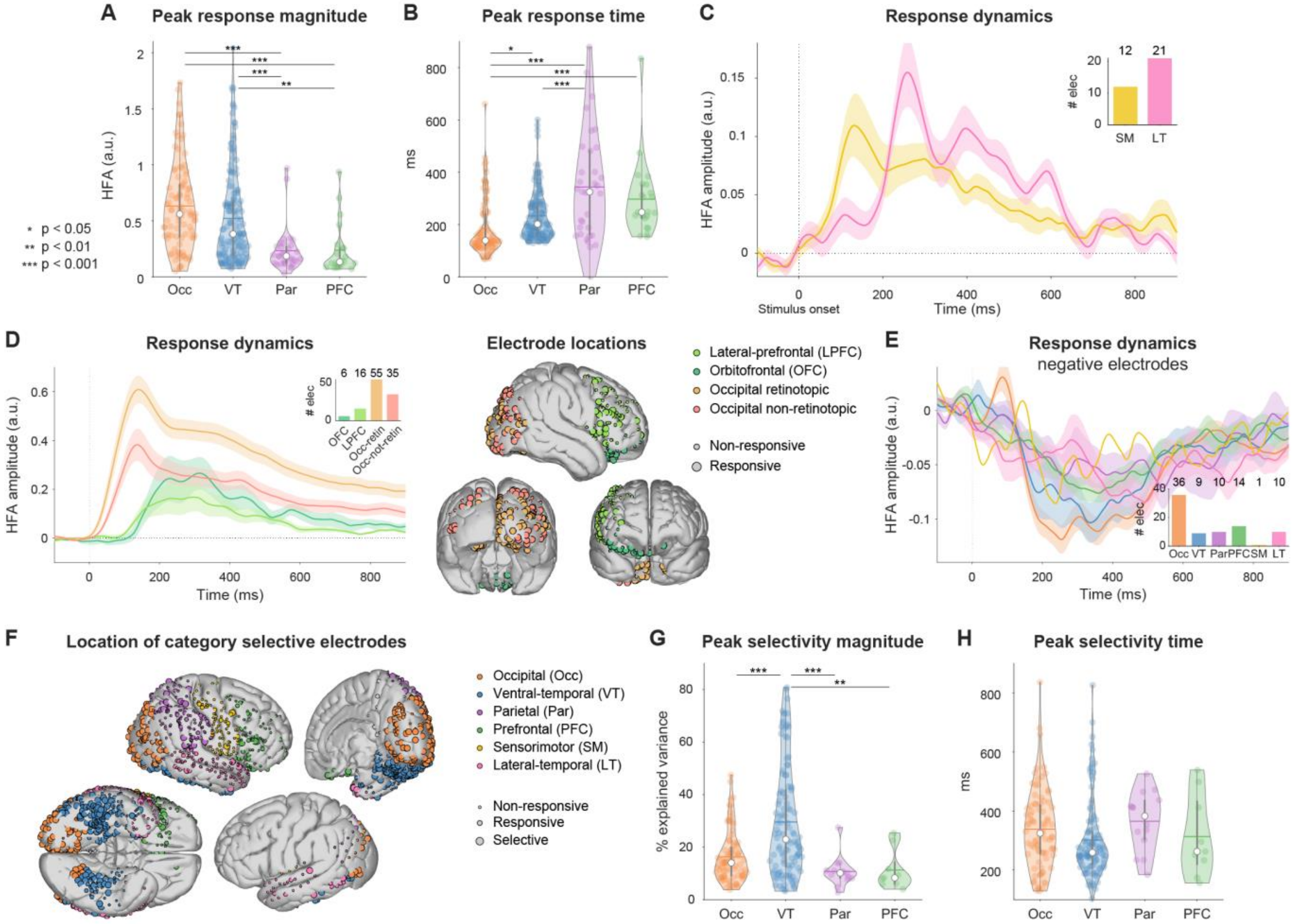


**Figure S1. Single electrode response properties.**

All images analyzed in the figure were presented for 900ms or longer. (**A-B**) Single electrode HFA response properties (corresponding to Figure 1D-E, only positively responding electrodes). (A) Peak HFA response was significantly higher in the classical visual regions (VT and Occ), relative to frontoparietal cortex (one-way ANOVA, $F(3,293) = 12.5$, $p < 10^{-6}$). Black horizontal lines connect regions which showed significant post-hoc differences (Tukey-Kramer method): all $p < 0.006$, all Cohen's $d > 0.7$. (B) Responses peaked first in Occ (mean ± SEM across electrodes: 187 ± 12ms), followed by VT (233 ± 8ms), PFC (297 ± 31ms) and Par (343 ± 36ms). Significant by one-way ANOVA ($F(3,293) = 15.73$, $p < 10^{-8}$). Post-hoc results shown in the figure: $p_{Occ\text{-}VT} < 0.019$, Cohen's $d = 0.47$, for all other significant comparisons: $p < 0.001$, $d > 0.9$. Similar results were found for half-peak times (not shown). (**C-E**) HFA response dynamics for regions or electrodes not shown in Figure 1D. For each electrode we averaged the trials from categories the electrode was responsive to; shaded area: SEM across electrodes. (C) SM and LT positively responding electrodes. (D) PFC subdivision: OFC and LPFC, and Occ subdivision: retinotopic and non-retinotopic regions. Dynamics plot: only positively responding electrodes included; electrode locations plot: all responsive electrodes. (E) Negatively responding electrodes from the six ROIs. No differences were observed in relative attenuation or latency between ROIs. (**F**) Location of category selective electrodes (by one-way ANOVA for four categories; including both positively and negatively responding sites). 236 electrodes were selective in at least one time-window (VT, 112; Occ, 83; Par, 14; PFC, 9; SM, 2; LT, 10). Single electrode selectivity was more prevalent during the onset response (100-300ms, 171 electrodes; 300-500ms, 193 electrodes; 500-700ms, 139 electrodes; 700-900ms, 100 electrodes). Interestingly, 16 electrodes began showing selectivity only after the first 500ms (6 VT, 6 Occ, 1 Par and 3 LT). (**G-H**) Univariate category selectivity properties corresponding to Figure 1F-G. Only selective electrodes are shown (as the goal was to estimate the dynamics of selective electrodes, not the absolute selectivity). (G) Peak selectivity magnitude was significantly higher in VT relative to all other regions (one-way ANOVA, $F(3,214) = 14.57$, $p < 10^{-7}$; post-hoc tests all $p < 0.008$, $d > 0.77$). (H) Time of peak selectivity did not differ between regions (one-way ANOVA, $F(3,214) = 1.82$, $p > 0.14$). Half-peak times were significantly different ($F(3,214)=4.82$, $p < 0.003$) with post-hoc tests showing selectivity in VT was faster relative to Par ($p < 0.008$, $d = 0.92$) and no other significant comparisons. Violin plots (A-B, G-H) were created using ‘violinplot.m’ in Matlab[1] – colored dots: single electrodes, colored horizontal lines: mean across electrodes, white dots: median across electrodes, gray vertical bars: interquartile range, contour lines: kernel probability density estimate (Matlab function ksdensity). Occ, occipital; VT, ventral-temporal; PFC, prefrontal; Par, parietal; SM, sensorimotor; LT, lateral-temporal; OFC, orbitofrontal; LPFC, lateral-prefrontal.

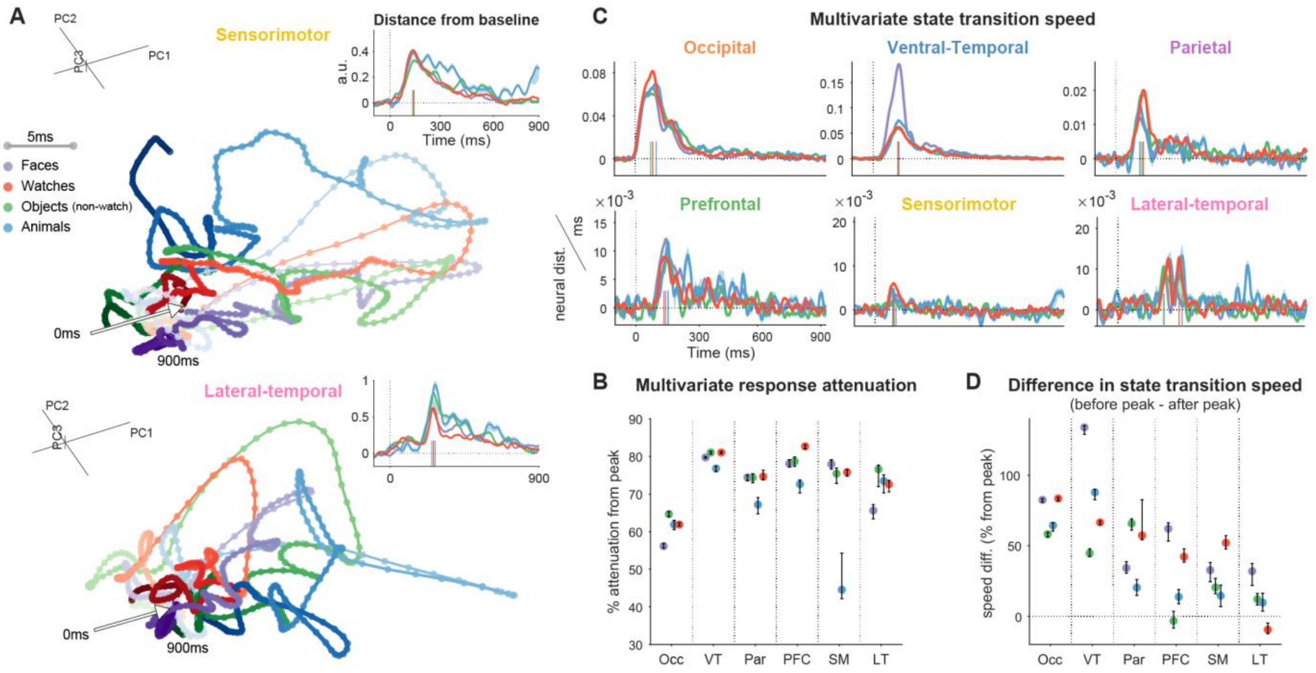


**Figure S2. State-space dynamics and state transition speed.**

All images analyzed in the figure were presented for 900ms or longer. (**A**) Multivariate state-space trajectories (projection on first 3 principal components; PCA using all responsive electrodes). Time is marked by the color and width of the trajectory (darker and thicker as the trial progresses). Dots are separated in time by 5ms. Responses to each category were averaged prior to PCA, which was used for visualization purposes only; characterization of response dynamics was carried out using the full multivariate response. Insets: point-by-point distance of each trajectory from the pre-stimulus state (computed using the full response, prior to PCA; baselined for presentation purposes). Dashed line: stimulus onset; colored vertical lines on the abscissa: peak distance times. (**B**) Multivariate response attenuation 800-900ms after onset relative to peak response. Most regions exhibited 70-80% attenuation (range across categories (%): VT: 77-81, Par: 67-75, PFC: 73-83, SM: 45-78, LT: 66-78), except for Occ (56-65%). Error bars denote 95% confidence interval (Jackknife estimate). Points correspond to categories (color code as in (A)). (**C**) State transition speed dynamics ("neural distance" travelled by the trajectory per ms) for the trajectories depicted in (A) (notations as in the inset). Transition from baseline to onset response state was very fast, reaching peak speed earliest in occipital cortex (73-100ms, across categories), followed by VT (117-120ms) and parietal cortex (113-130ms) and slowest in PFC (133-152ms). No comparable peak was seen for the return direction, from active state to the pre-stimulus state. (**D**) Difference between state transition speed before and after peak response magnitude, defined as the time of peak distance from baseline (insets in (A)). In all but two cases ($PFC_{objects}$, $LT_{watches}$) the speed before the peak (state transition from baseline to active) was higher than the return direction. This was especially prominent in VT and Occ, which settle into sustained activity states tracking the duration of the stimulus (see Figure S3). We used the mean state transition speed 100ms before\after peak response (similar results were obtained using different time ranges). Values were scaled by the instantaneous speed at the peak response to enable comparison between regions and categories. Error bars and color code as in (B).

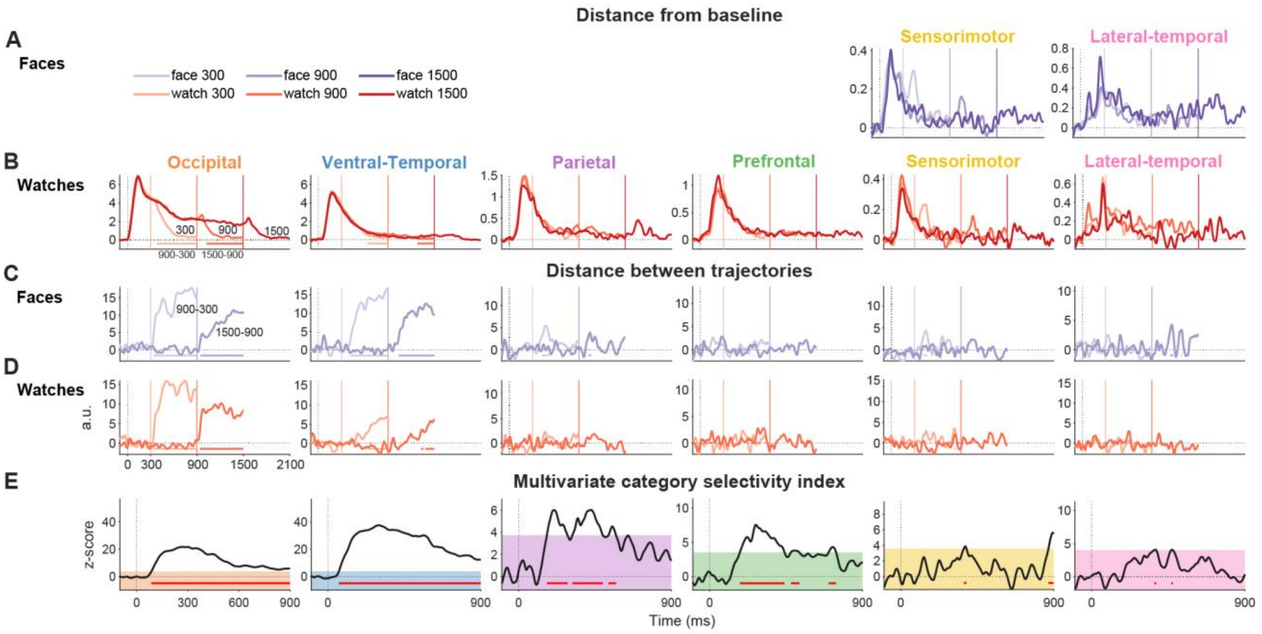


**Figure S3. Multivariate responses in sensory regions track the duration of the stimulus.**
(**A-B**) Multivariate trajectory distance from the baseline state per duration (baselined for presentation purposes; darker lines correspond to longer stimuli; stimuli offsets are marked by a vertical line of the same color). (A) Faces: SM and LT regions, complementing Figure 2B. (B) Watches (all ROIs). Response dynamics tracked the duration of the stimulus only in Occ and VT. Trajectories are cropped 600ms after stimulus offset corresponding to the shortest ISI (beyond that point another stimulus could be presented). Horizontal bars: significant differences between durations (permutation, max-statistic control for multiple comparisons, $p < 0.05$). Bar color corresponds to the shorter duration in the comparison (bright: 900-300ms, dark: 1500-900ms). Distance magnitude is comparable in the same ROI between time-points, but not between ROIs, as it also depends on the number of electrodes in the region. (**C-D**) Point-by-point distances between trajectories (without considering the baseline state; 900-300ms, lighter color and 1500-900ms, darker color). (C) Faces. (D) Watches. Traces were z-scored on a time-point by time-point basis using a null distribution obtained from shuffling duration labels. Only in Occ and VT response trajectories tracked the duration of the stimulus. Transient differences were observed in Par, SM and LT, possibly reflecting offset responses. PFC responses did not show a similar trend. Notations as in (A-B). (**E**) Category selectivity index (measuring the dispersion between responses to different categories, see Methods for more details) for stimuli with duration of 900ms or longer. All regions showed significant category information, but it was sustained only in sensory regions (Occ, 87-900ms; VT, 65-900ms; Par, 167-288ms, 317-496ms and 530-573ms; PFC, 178-440ms, 479-528ms and 701-744ms; SM, 372-386ms, 870-900ms; LT, 368-379ms, 468-476ms). Horizontal red bars: above chance selectivity using permutation max-statistic control for multiple comparisons ($p < 0.05$). Shaded regions: 95% percentile of the null distribution.

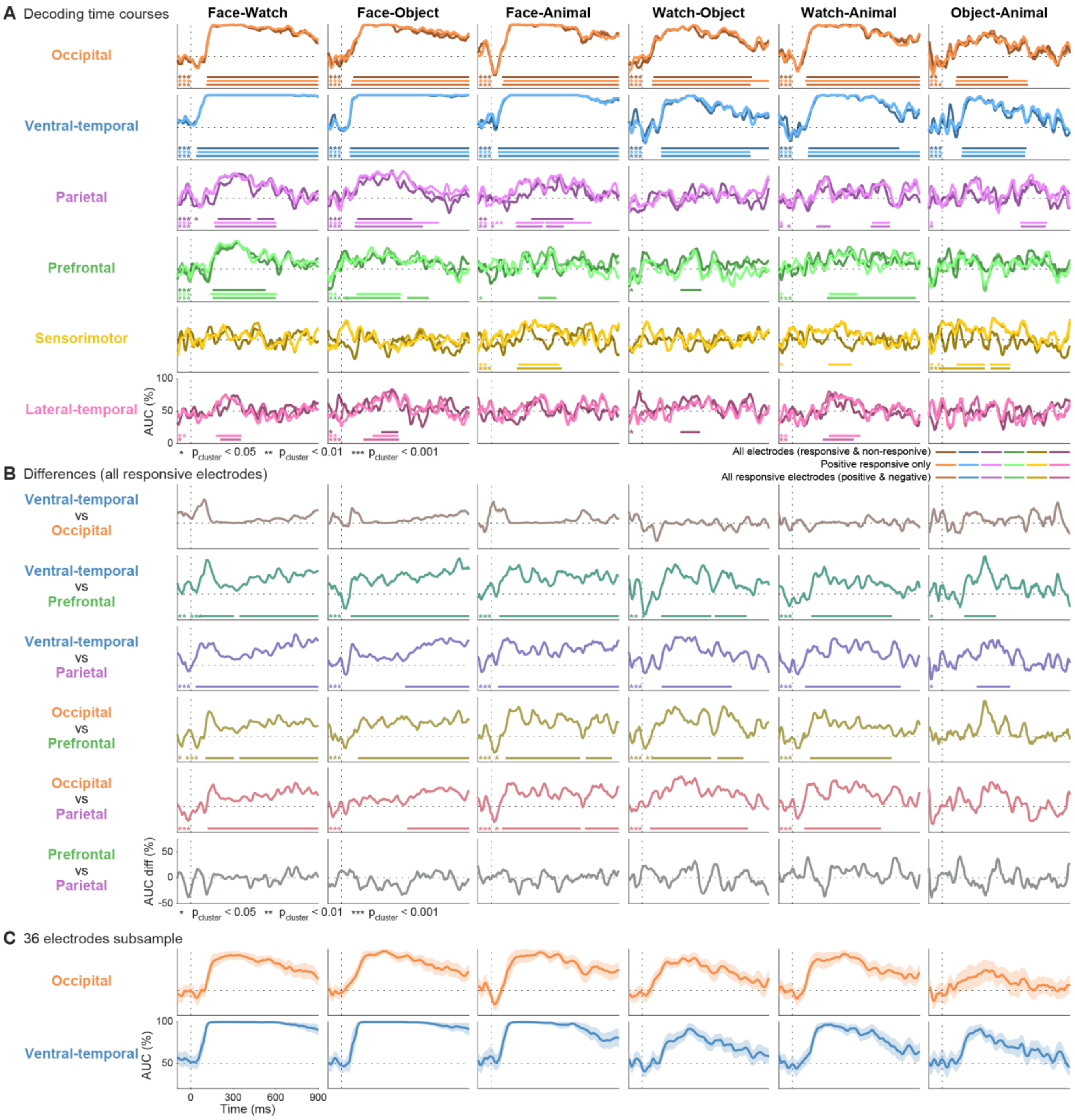


**Figure S4. Sustained coding in sensory regions and transient coding in frontoparietal regions.** All panels depict decoding performance (AUC) for images presented for 900ms or longer. Columns: category comparisons. (**A**) Decoding dynamics for all pairwise comparisons in all cortical ROIs. Color-coding according to the electrodes used for the analysis (Table S4): Darkest lines – all electrodes (responsive and non-responsive), intermediate shading – all responsive electrodes (corresponding to the main figures), brightest lines – decoding using only positively responding electrodes (responsive electrodes which increased their activity relative to baseline). Dashed lines: stimulus onset and chance level (AUC = 50%). Horizontal bars: significant clusters (cluster-based permutations); cluster p-values are indicated by the asterisks at the bottom-left corner (corresponding to the cluster temporal order). Occ and VT: All pairwise comparisons were significant, with clusters emerging shortly after stimulus onset and persisting throughout stimulus presentation for all comparisons except animal-object, possibly due to reduced power as these two categories had the smallest number of exemplars. Par and PFC also showed reliable category information. Using all responsive electrodes, in both regions we found faces were significantly distinguishable from all other categories, and animals were distinguishable from watches. $Par_{animal\text{-}object}$ was significant using cluster permutations, but not significant considering only the peak response (Figure 3B), indicating a weak and broad effect, while $Par_{watch\text{-}object}$ was significant using max-statistic for the peak response but not with cluster permutations, indicating a strong yet brief category signal. Considering only positively responding electrodes led to similar results, except the $PFC_{face\text{-}animal}$ was not significant. Adding the non-responsive electrodes led to poorer decoding since it introduced substantial noise; nonetheless, both regions showed significant category information. SM showed sensitivity to the animal category and LT showed weak animacy information. $LT_{face\text{-}watch}$, $LT_{face\text{-}object}$ and $LT_{watch\text{-}animal}$ were all significant ($LT_{object\text{-}animal}$ was not, but this comparison had lower power). (**B**) Pairwise contrasts of decoding time-courses between the four core ROIs. Lines mark the difference between decoding time-courses. Notations and significance as in (A). ROIs can be grouped into two distinct groups, based on the temporal dynamics of decoding: VT and Occ show sustained coding, while PFC and Par show transient coding. Differences within each group were not significant (VT-Occ and PFC-Par show no significant differences), and differences between groups were all significant except $Occ\text{-}PFC_{animal\text{-}object}$ and $Occ\text{-}Par_{animal\text{-}object}$. (**C**) Decoding in VT and Occ using only 36 responsive electrodes (equivalent to the number of responsive electrodes in PFC). Even after reducing the number of electrodes category information remained high throughout stimulus presentation (especially in VT, compare to panel (A)). We repeated the procedure $N_{iterations}$ = 1,000, each time using a different electrode subset. Lines and shaded area: mean ± 1 standard deviation across iterations.

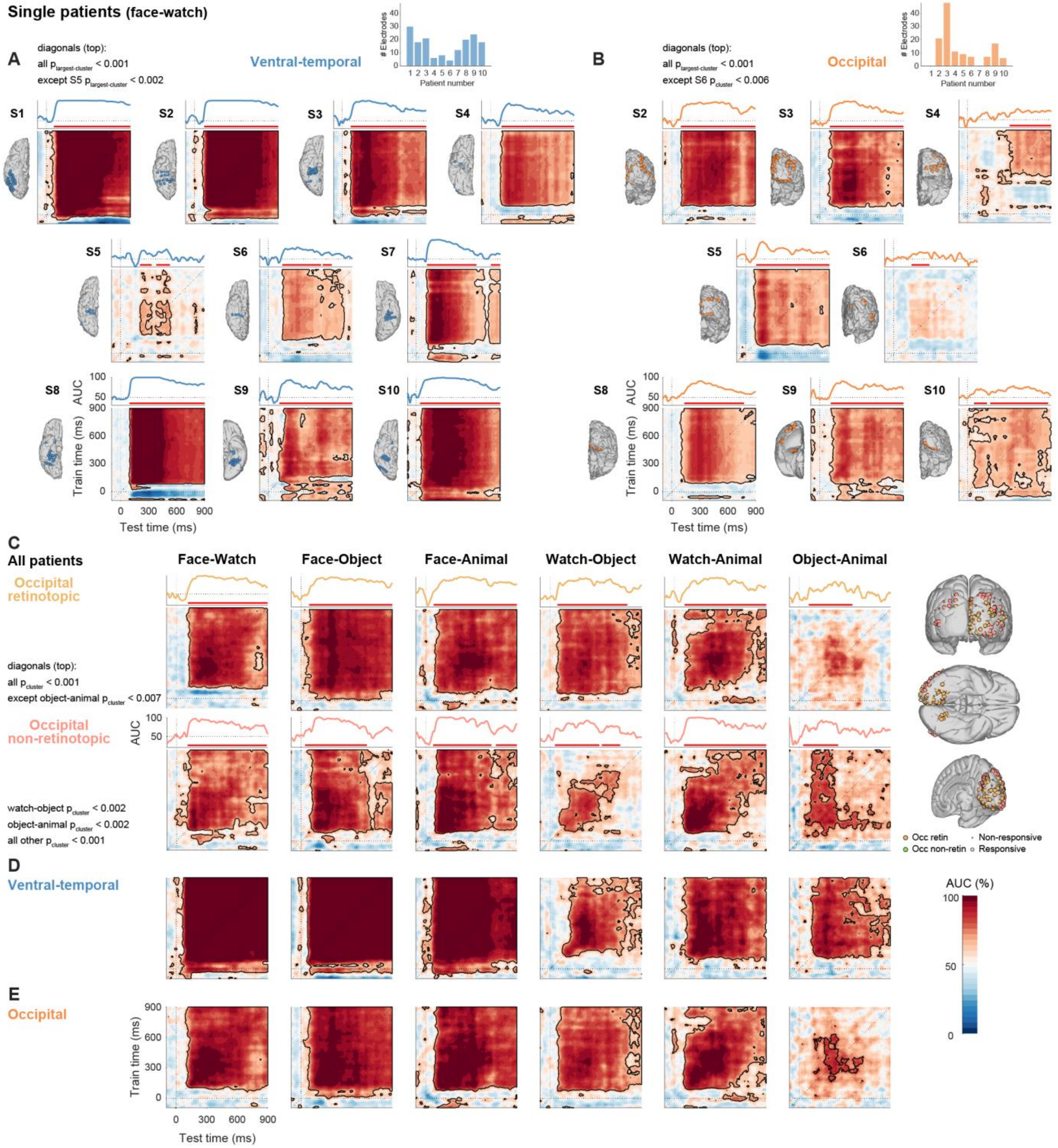


**Figure S5. Visual sensory regions maintain a sustained and stable representation of category information for multiple categories and at the single patient level.**

All panels depict decoding (AUC) for images presented for 900ms or longer. (**A-B**) Single patient results (face-watch comparison) for all patients with responsive electrodes in VT (10/10 patients) and Occ (8/10 patients). Each panel shows the temporal generalization matrix (TGM) of a single patient. Contiguous significant points contoured in black ($q_{FDR} < 0.05$, permutation test FDR corrected). Line plots above each TGM depict the matrix diagonal (training and testing on the same time-point), with statistical significance (cluster-based permutations) marked by red horizontal bars. Dashed lines: stimulus onset and chance level (AUC = 50%). Bar plots on the top right of each panel show the number of electrodes in each patient. Electrode positions on the native brain are depicted next to each panel. In both regions, the majority of patients showed high decoding which was sustained and stable throughout stimulus duration (all $p_{largest\text{-}cluster} < 0.001$ except $VT_{S5}$ $p_{largest\text{-}cluster} < 0.002$ and $Occ_{S6}$ $p_{cluster} < 0.006$). Differences between patients likely stem from the variability in electrode coverage, as peak AUC, mean AUC and mean generalization (off-diagonal decoding; both means after 100ms) were correlated across patients with the number of responsive electrodes in the region (Spearman correlations, one-sided tests). VT: $\rho_{peak} = 0.66$, $p < 0.019$; $\rho_{on\text{-}diag} = 0.72$, $p < 0.009$; $\rho_{off\text{-}diag} = 0.52$, $p < 0.061$. Occ: $\rho_{peak} = 0.65$, $p < 0.046$; $\rho_{on\text{-}diag} = 0.72$, $p < 0.026$; $\rho_{off\text{-}diag} = 0.72$, $p < 0.026$. (**C**) Across patients decoding for all comparisons separately for Occ retinotopic and non-retinotopic regions, electrode locations shown on the right. Notations as in (A-B). Decoding in both subregions was high (peak AUC (mean ± SEM across comparisons): retinotopic, 97 ± 2%, non-retinotopic, 97 ± 1.6%; all $p_{cluster} < 0.007$). All category comparisons yielded significant clusters in both regions (diagonals (top): retinotopic, all $p_{cluster} < 0.001$ except object-animal $p_{cluster} < 0.007$; non-retinotopic, all $p_{cluster} < 0.001$ except watch-object, object-animal $p_{cluster} < 0.002$), yet both subregions showed sustained and stable decoding for some, though not all, comparisons (mean AUC after 100ms: retinotopic, 83 ± 4.1%, non-retinotopic, 79.1 ± 2.9%; mean generalization (off-diagonal AUC) after 100ms: retinotopic, 78.7 ± 4.4%, non-retinotopic, 73.5 ± 2.9%). Object-animal decoding was low in both subregions, possibly because of the large heterogeneity in low-level features, though the comparison may be underpowered relative to the others since these two categories contained the lowest number of exemplars. (**D-E**) VT and Occ TGMs for all pairwise comparisons (complementing Figure 3D; see Figure S4A-B for the diagonals). Notations as in the previous panels. Both regions show highly stable coding of most category comparisons, indicated by the rectangular shape of the TGMs.

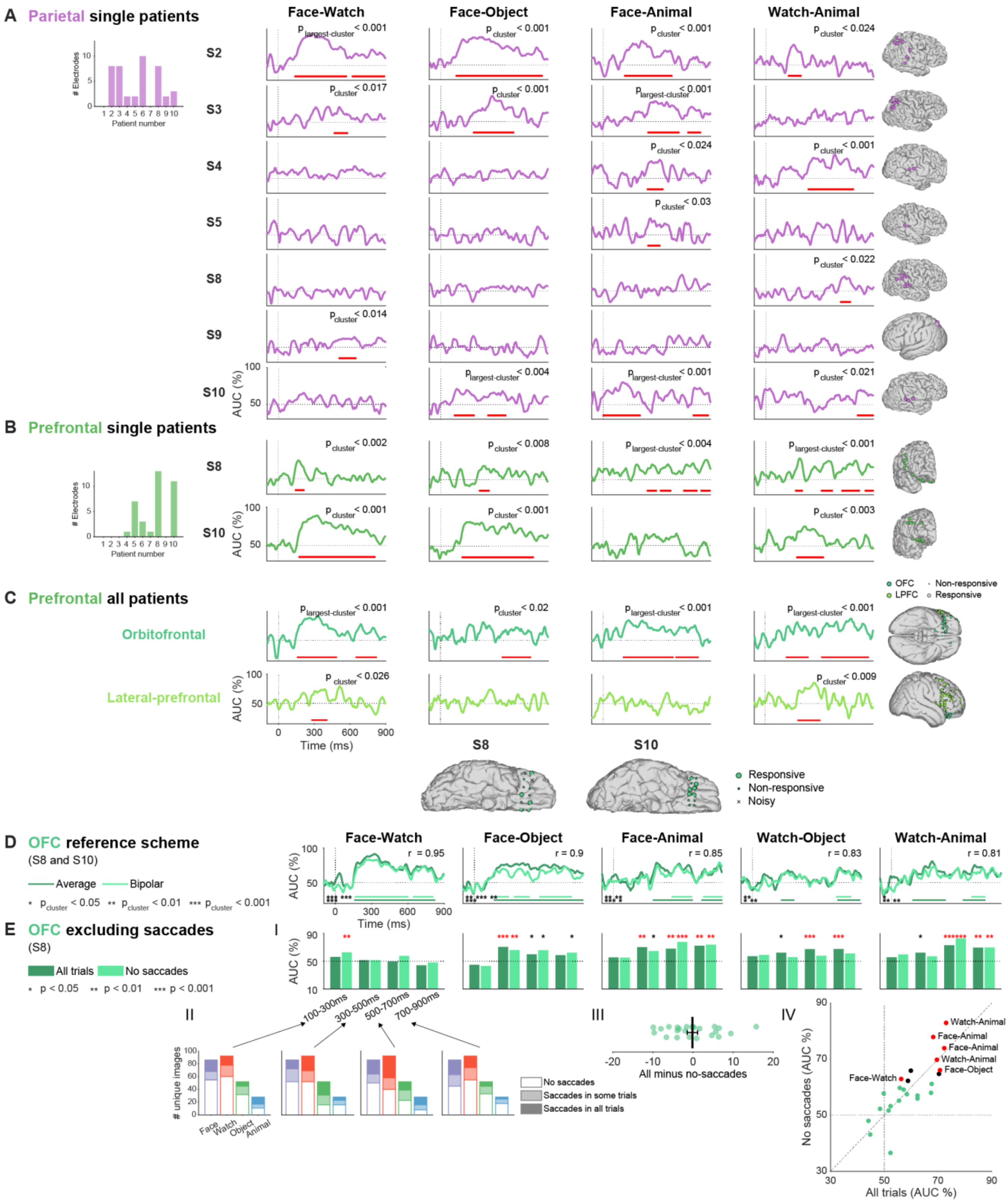


**Figure S6. Visual category information in prefrontal and parietal cortex is found in multiple patients and is not the result of saccadic events.**

All stimuli presented for 900ms or longer. (**A-B**) Single patient decoding performance in Par and PFC for the four comparisons which were significant in both regions (Figure S4C-D). Electrode positions on the native brain are depicted on the right. Horizontal red bars: significant clusters (cluster-based permutations; cluster p-values shown in the top-right corner). Dashed lines: stimulus onset and chance level (AUC = 50%). (A) Par responsive electrodes were present in eight patients (bar plot inset), seven of which are plotted (S6 did not show significant decoding for the plotted comparisons). In addition to the plotted time-courses, the following comparisons were also significant: $S6_{object\text{-}animal}$ (cluster time-range: 635-770ms, $p_{cluster} < 0.02$) and $S4_{watch\text{-}object}$ (450-715ms, $p_{cluster} < 0.002$). (B) PFC responsive electrodes were present in six patients (bar plot inset). Only patients with more than 10 responsive electrodes are plotted (S8: 8 OFC, 5 LPFC; S10: 5 OFC, 6 LPFC). Other than the plotted comparisons, object-watch decoding was significant in both patients (S8: 160-205ms, 320-405ms, 420-495ms, 515-570ms, 700-755ms and 800-875ms, $p_{largest\text{-}cluster} < 0.014$; S10: 265-410ms, $p_{cluster} < 0.011$). Other patients did not show significant decoding as their prefrontal coverage was very partial, with either a single (2 patients), three (1 patient) or seven responsive electrodes (1 patient). (**C**) PFC decoding performance separately for OFC and LPFC (all patients, 22 and 14 responsive electrodes respectively, shown on the right). Only comparisons which PFC was sensitive to are plotted (watch-object and object-animal decoding were not significant in both sub-regions). Decoding was significant in both regions, though it was higher in OFC (peak AUC (mean ± SEM across six category comparisons): OFC, 83.7 ± 2%; LPFC, 77.3 ± 2.3%). Notations as in (A-B). (**D-E**) OFC category information was not driven by eye-muscle activity (see Methods for more details). (D) We repeated the decoding analysis in OFC using a bipolar montage, which emphasizes local neural activity as opposed to more distant signals (such as ocular contamination). Category information remained significantly decodable, and decoding time-courses were highly correlated between the montages (top-right corner; mean ± SEM across category comparisons: 0.83 ± 0.04). Object-animal not shown since it was not significant for either reference scheme (r = 0.67). Notations as in (A-C); dark green: average reference, Bright green: bipolar montage; significance of each time-course marked by bars of the same color, cluster p-values indicated by the asterisks at the bottom-left corner (corresponding to the cluster temporal order). The slight differences between average reference decoding in (D) and OFC decoding in (C) stems from the electrodes used in each analysis: panel (C) uses all responsive OFC electrodes from all patients, while (D) uses only electrodes from grids located fully over PFC (see Methods). (E) Category information was present in OFC in trials without any saccades and was highly consistent with decoding performance using all trials. Analysis performed on data from one patient (S8) where we were able to identify the timing of saccades reliably (Methods). (I) Analysis was performed separately in four non-overlapping time-windows from 100 to 900ms, once including all trials (dark bars), and once only using trials where no saccadic events occurred in this time-range (bright bars). Despite the reduction in the number of exemplars, decoding in most comparisons survived (and new comparisons gained significance). Statistical testing was performed separately for each comparison, time-window and trial group using permutation testing (one-sided); p-value (uncorrected) is marked by the number of asterisks, comparisons surviving Bonferroni correction for multiple time-windows are marked in red. (II) The number of unique images in each category and each time-window, split into: images where all trials did not contain saccades in this time-window (white fill), images where some trials included saccades (bright fill, trials with saccades were excluded from the analysis without saccadic events, other trials of the same image were included) and images for which all trials included saccades (dark fill, excluded in the analysis without saccadic events). Across categories and time-windows 11–43% of images were excluded due to the presence of saccades (mean ± SEM: 24 ± 3%). (III) Difference in decoding with all trials versus decoding without saccadic events was not significantly different from zero (mean ± SEM across categories and time-windows: 0.1 ± 1.3%). Each dot stands for decoding results in a different comparison and different time window. Vertical black line: mean, error bars: SEM. (IV) Results were highly correlated between analysis conditions ($r(22) = 0.8$, $p < 0.0001$). Each dot stands for decoding results in a different comparison and different time window (x-axis: all trials, y-axis: only trials without any saccadic events). Dashed lines: chance level (AUC = 50%) and equality (x=y). Comparisons which were significant ($p_{perm} < 0.05$) in the analysis which did not include any saccadic events are marked in red (Bonferroni corrected for four windows) or black (significant only uncorrected, corresponding to the single stars in (I)). Altogether, the presence of saccades did not increase the likelihood of successful decoding.

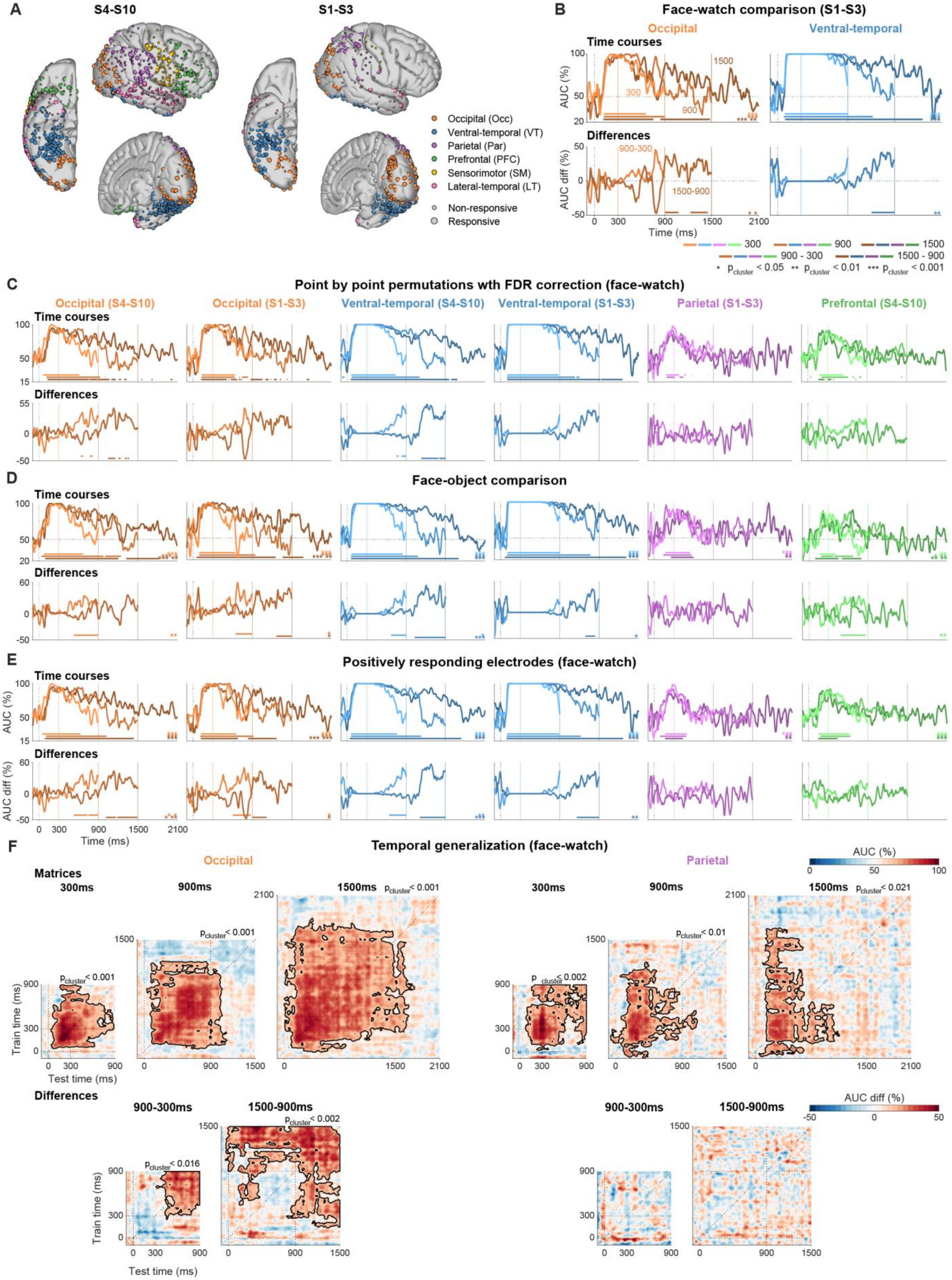
A
S4-S10
S1-S3
Occipital (Occ)
Ventral-temporal (VT)
Parietal (Par)
Prefrontal (PFC)
Sensorimotor (SM)
Lateral-temporal (LT)
Non-responsive
Responsive
B
Face-watch comparison (S1-S3)
Occipital
Ventral-temporal
Time courses
Differences
AUC (%)
AUC diff (%)
Time (ms)
300
900
1500
900 - 300
1500 - 900
* $p_{cluster} < 0.05$ ** $p_{cluster} < 0.01$ *** $p_{cluster} < 0.001$
C
Point by point permutations wth FDR correction (face-watch)
Occipital (S4-S10)
Occipital (S1-S3)
Ventral-temporal (S4-S10)
Ventral-temporal (S1-S3)
Parietal (S1-S3)
Prefrontal (S4-S10)
D
Face-object comparison
E
Positively responding electrodes (face-watch)
F
Temporal generalization (face-watch)
Occipital
Parietal
Matrices
300ms
900ms
1500ms
Train time (ms)
Test time (ms)
AUC (%)
AUC diff (%)
Differences
900-300ms
1500-900ms
$p_{cluster} < 0.001$
$p_{cluster} < 0.002$
$p_{cluster} < 0.01$
$p_{cluster} < 0.021$
$p_{cluster} < 0.016$

**Figure S7. Decoding in sensory regions, but not frontoparietal regions corresponds to the duration of experience.**

All panels except for (D) depict decoding of faces versus watches. (**A**) Electrode positions for patients S4-S10 and patients S1-S3 (see Methods for the rationale behind this split). Left hemisphere electrodes were projected to the right hemisphere. For the number of exemplars viewed in each duration by each patient group see Table S6; the number of responsive electrodes for each patient group in each ROI is shown in Table S7. (**B**) Occ and VT from patients S1-S3 (complementing Figure 4; not shown: $PFC_{S1\text{-}S3}$ since S1-S3 did not contain any responsive PFC electrodes, $Par_{S4\text{-}S10}$ was not significant). Decoding time-courses in both regions corresponded to the duration of the stimulus. Both showed prolonged onset responses, eliminating differences between 300 and 900ms stimuli, but the difference between 1500 and 900ms stimuli was significant. Top: decoding performance for each stimulus duration (darker lines correspond to longer stimuli, offsets marked by vertical lines of the same color), bottom: difference between durations (1500 vs 900ms, dark; 900 vs 300ms, bright). Results are displayed from 100ms before onset to 600ms after offset of the shortest stimulus, corresponding to the minimal ISI in the task as beyond that point another stimulus could be presented. Dashed lines: stimulus onset and chance level (AUC = %50). Horizontal bars: significant clusters (cluster-based permutations); cluster p-values are indicated by the asterisks at the bottom-right corner (corresponding to the cluster temporal order). In the difference plots only time-points after the offset of the shorter stimuli were considered (results are highly similar considering the entire presented range). (**C**) Statistical testing using point-by-point permutation testing with FDR correction ($q_{FDR} < 0.05$) corresponding to (B) and Figure 4A-B. Notations as in (B). (**D**) Decoding faces versus objects – similarly to the face-watch decoding results, decoding in visual regions, but not frontoparietal regions tracked the duration of the stimulus. Other category comparisons were not possible due to paucity of exemplars in the animal category (Table S6). For patients S1-S3 we used four-fold cross validation in this analysis to ensure at least two exemplars of each category were present in each fold. (**E**) Decoding using only electrodes which increased activity relative to the pre-stimulus state. (D-E) Notations and significance as in (B). (**F**) Temporal generalization matrices for face-watch decoding in each stimulus duration (top) and the differences between durations (bottom) in $Occ_{S4\text{-}S10}$ and $Par_{S1\text{-}S3}$ (complementing Figure 4C-D). Dashed lines: stimulus onset, offset, and the diagonal. Significant clusters are contoured in black; cluster p-values are shown in the top-right corner.

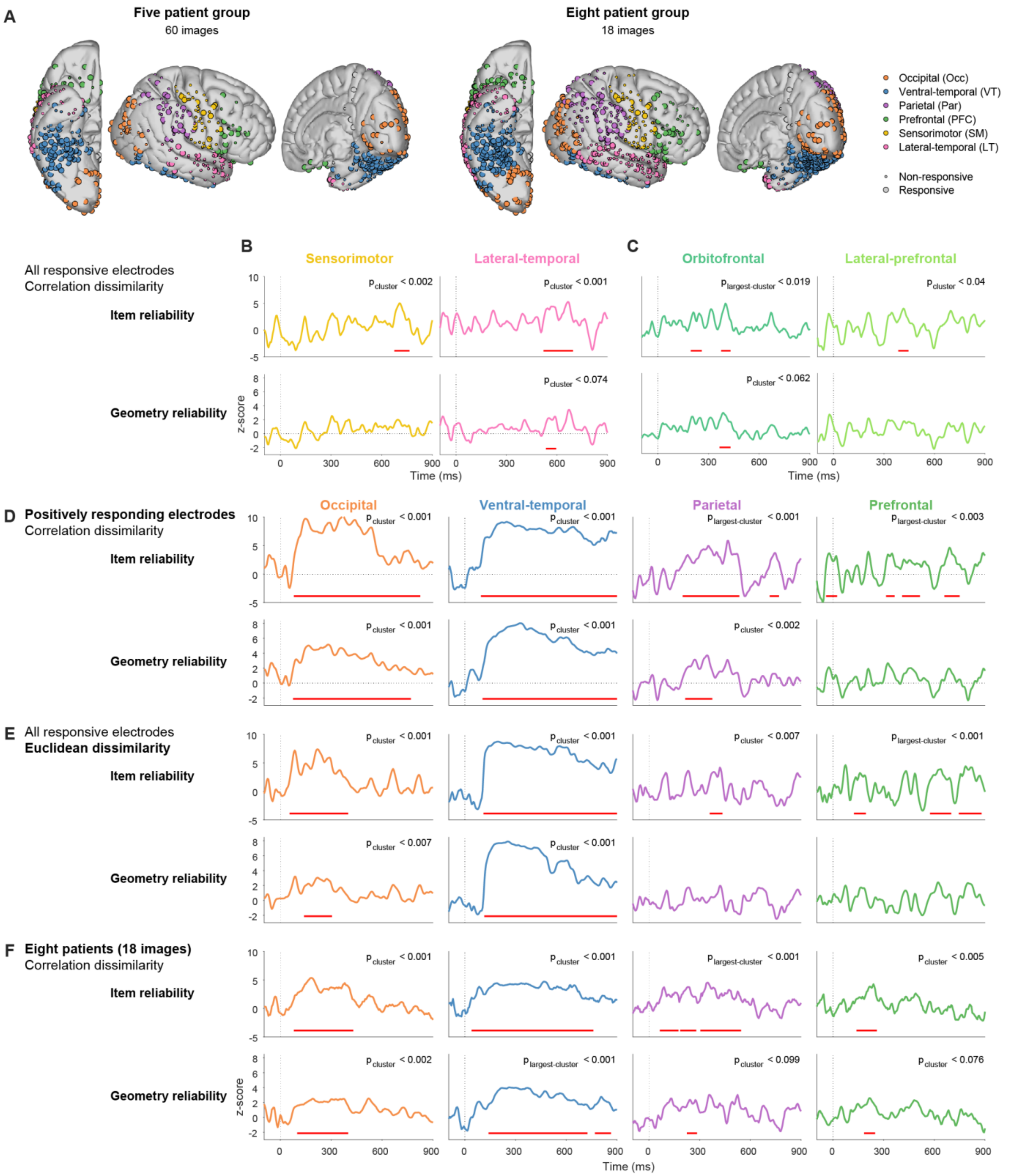


**Figure S8. Exemplar representation is sustained in visual regions, especially ventral temporal cortex and transient in frontoparietal regions.**

All images analyzed in the figure were presented at least twice for 900ms or longer. (**A**) Electrode locations for the five-patient group (S1, S2, S5, S7, S8; 60 images) and the larger eight-patient group (adding patients S6, S9, S10;

18 images). Left hemisphere electrodes were projected to the right hemisphere. The number of responsive electrodes for each subgroup in each ROI is shown in Table S7. (**B-F**) Item Reliability (IR, top) and Geometry Reliability (GR, bottom) dynamics. Dashed lines: stimulus onset and chance level (no single-item information). Red horizontal bars: significant clusters (cluster-based permutations); the corresponding cluster p-value is shown in the top-right corner. (B-C) Correlation dissimilarity using all responsive electrodes of the main five-patient group (regions (B) and sub-regions (C) not shown in Figure 5D-E). (D-F) IR and GR in the main four ROIs (complementing Figure 5D-E): (D) Using only positively responding electrodes (correlation dissimilarity, five-patient group); (E) Using Euclidean dissimilarity (all responsive electrodes, five-patient group); (F) Using all responsive electrodes from the eight-patient group (correlation dissimilarity). All approaches converged to the same conclusion – single item information is reliably maintained in VT and Occ, though it is more sustained in VT. Par and PFC both show significant IR, yet inconsistent GR, suggesting that only part of the representational geometry is preserved.

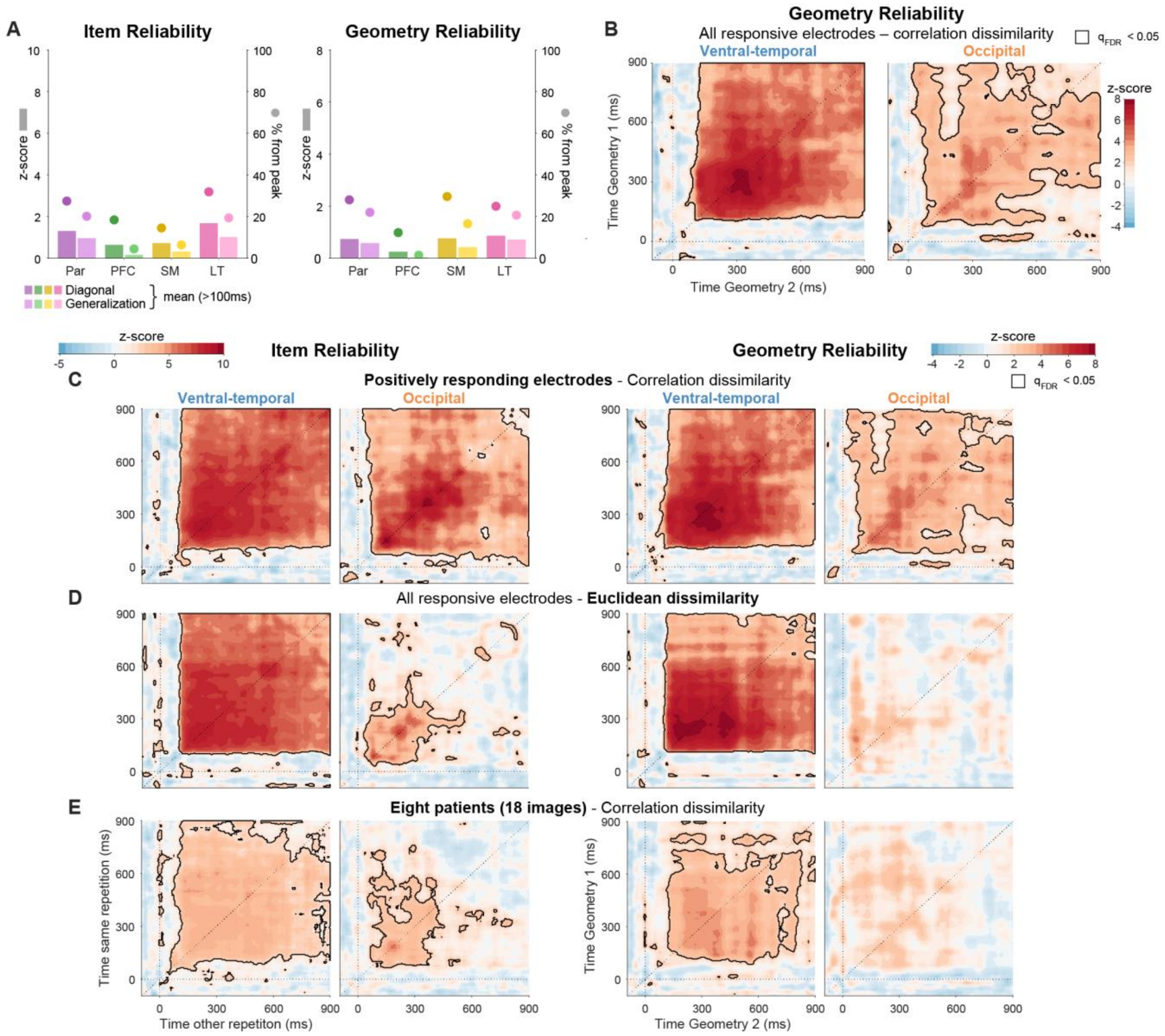


**Figure S9. Exemplar-level information is highly stable in ventral temporal cortex, and less robust in occipital cortex.**

All images analyzed in the figure were presented at least twice for 900ms or longer. (**A**) Representation reliability within the same time-point (diagonal) and stability of representation between time-points (across repetitions; generalization\off-diagonal) for regions not shown in Figures 5F and S9B. Left: Item Reliability (IR, item-by-item position within the neural representation, Figure 5A), right: Geometry Reliability (GR, reliability of the full representational geometry, Figure 5B). Bars: mean value relative to the null distribution (z-score units; left y-axis), dots: relative value (% from peak; right y-axis). (**B**) GR stability between time-points in VT and Occ (complementing IR stability matrices in Figure 5F). Diagonals (dashed lines) correspond to the dynamics time-courses in Figure 7E. Significance using permutation testing with FDR correction for multiple comparisons (contiguous points with $q_{FDR} < 0.05$ are contoured in black). (**C**-**E**) Exemplar representation stability across time-points in VT and Occ (complementing Figure S8D-F), showing robust and stable exemplar representation in VT, but less so in Occ which does not persist with Euclidean dissimilarity or the larger patient group. Left: IR stability, right: GR stability. Statistical significance as in (B). (C) Only positively responding electrodes (correlation dissimilarity; five-patient group). (D) Euclidean dissimilarity (all responsive electrodes; five-patient group). (E) Eight-patient group (correlation dissimilarity, all responsive electrodes).

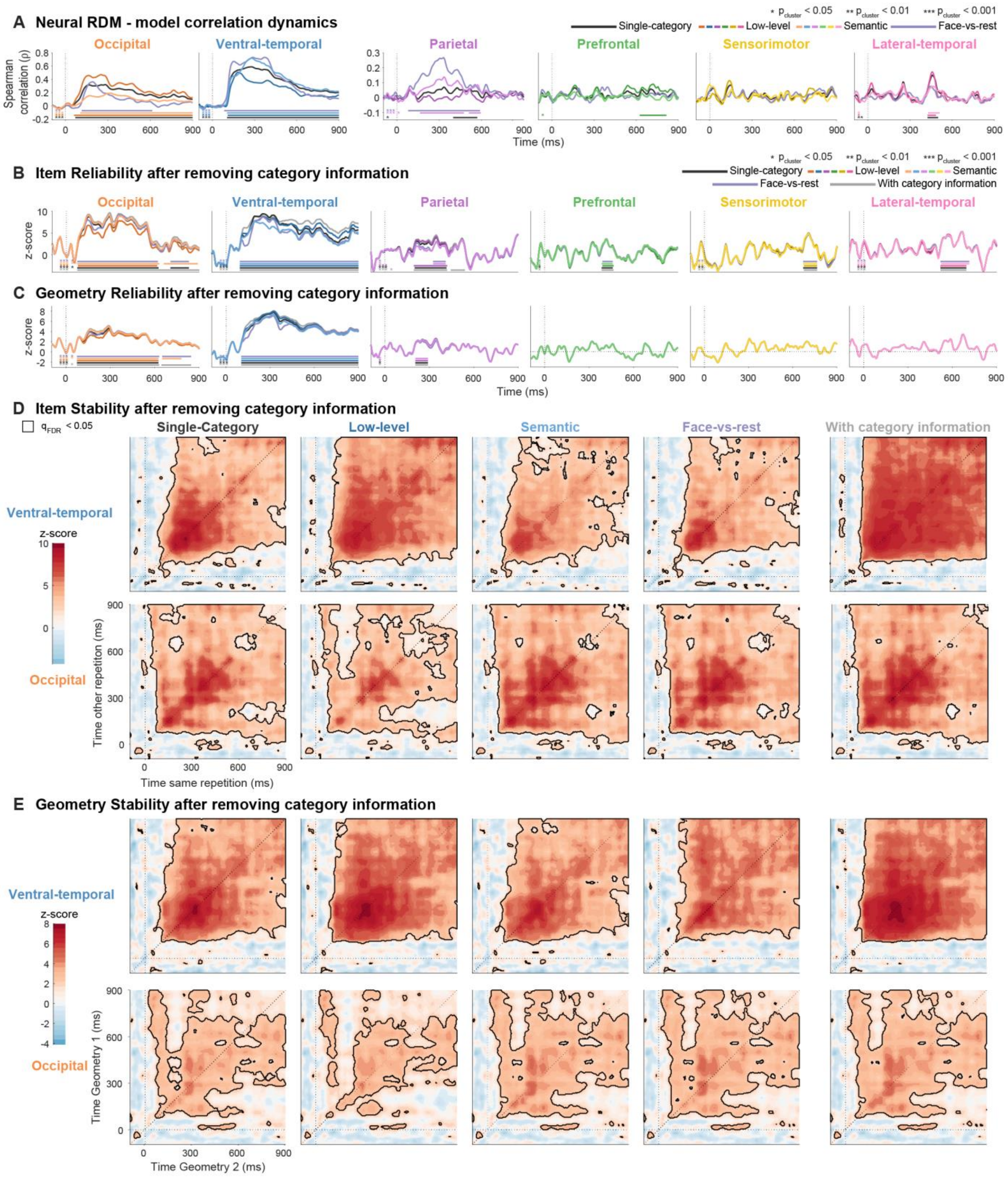


**Figure S10. Stability of the neural geometry is not fully accountable by category structure.** All images analyzed in the figure were presented at least twice for 900ms or longer. (**A**) Dynamics of Spearman correlation between the neural RDM (mean of the four RDMs in Figure 5B) in each ROI and each of the category models (for the model details see Figure 5C and Methods). Note the scale difference between Occ and VT to the other ROIs. Unsurprisingly, Occ was most correlated with the "Low-level" model throughout stimulus presentation ($\rho_{peak}$ = 0.47). VT was most correlated with the "Face-vs-rest" model ($\rho_{peak}$ = 0.73) until approximately 430ms after stimulus onset, and from that point with the "Semantic" model ($\rho_{peak}$ = 0.72). Par was most correlated with the "Face-vs-rest" model ($\rho_{peak}$ = 0.27). These high correlations reveal the neural RDMs contain category information, however, it also shows that a substantial proportion of the information within each RDM is not captured by categorical structure (mean unexplained variance ($1-\rho^2$) after 100ms: Occ, 91%; VT, 78%; Par, 98%). We asked whether this unexplained information is reliable and stable. Other regions showed only brief correlations with category models, therefore single-item information found in these regions is unlikely to be attributed to category affiliation, but for completeness we include them in the following analyses as well. In PFC, the difference between the low correlation with category models and our decoding results showing category information (Figure 3) is likely due to the partial coverage, stemming from the exclusion of patients who did not view enough exemplars twice (Methods). Lines are color-coded according to the correlated category model. Across panels, colors are consistent for "Single-category" (black) and "Face-vs-rest" (purple). "Low-level" and "Semantic" colors are based on the ROI color (darker and brighter shade respectively). Significant clusters (cluster permutation analysis) of above chance correlation are marked by horizontal bars of the same color. Cluster p-value are indicated by the asterisks at the bottom left side of each plot (corresponding to the cluster temporal order). Dashed lines: stimulus onset and chance level (no correlation). (**B-C**) Item Reliability (IR) and Geometry Reliability (GR) after partialling-out category information as captured by each of the models. Lines are color-coded according to the partialled-out model (color-code as in (A)). For comparison, gray lines depict the original IR\GR values, computed without partialling-out information (Figure 5D-E). Significance notations as in (A). Both measures were attenuated by the removal of category information but remained significant in all regions, indicating that exemplar-level information is maintained over and above the category distinctions. Occ reliability was no longer sustained after removing some of the category models, and after removing the "Face-vs-rest" model GR was longer significant in Par. (**D-E**) IR and GR stability (generalization in time) after removing category information in each model, showing sustained and stable reliability in VT, even after removing category information. In Occ exemplar representation stability was less robust to removal of category information (especially GR stability). Diagonals correspond to the reliability time-courses in (B-C). Stability without removing category information is shown in the right column for comparison (identical to Figure 5F (IR) and Figure S9B (GR)). Statistical significance using permutation testing with FDR correction (contiguous points with $q_{FDR} < 0.05$ are contoured in black).

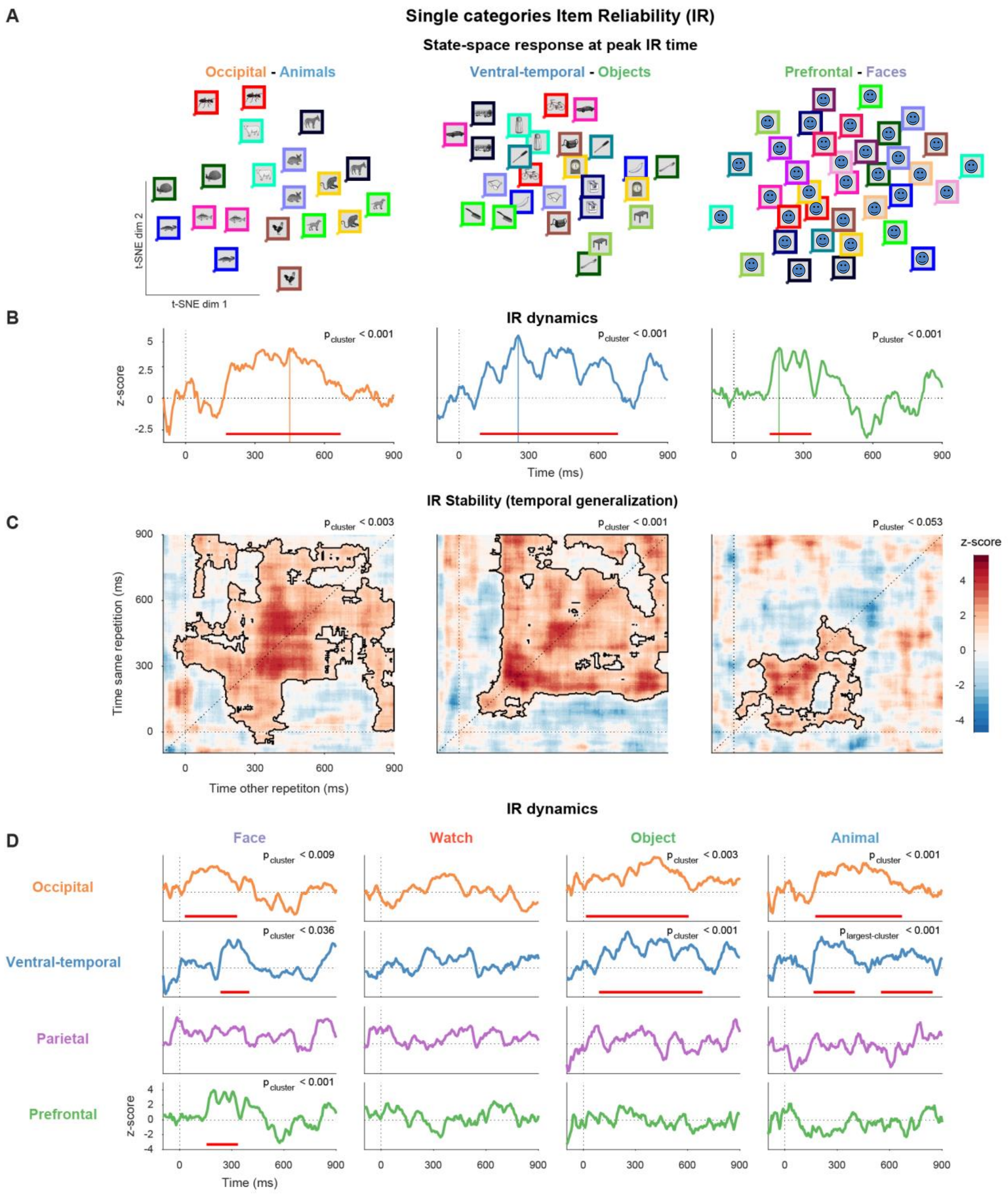


**Figure S11. Exemplar-level information is present within single categories.**
All images presented for at least twice for 900ms or longer. To account for the substantial reduction in the number of exemplars (between 3-fold and 6-fold reduction, depending on the category), data was smoothed by a 100ms moving window prior to reliability calculations. (**A**) Visualization of neural population responses to two presentations of all images from one example category, during the time of peak Item Reliability (IR; Occ-animals, 450ms; VT-objects, 255ms; PFC-faces, 195ms). Repetitions of the same image were represented more closely than presentation of different exemplars, indicating reliable exemplar representation within single categories. Image

frames are color-coded according to exemplar identity to ease comparison between repetitions (not presented during the task; color-code generated using ‘distinguishable_colors’[2]). The location of each image corresponds to the population response to that image, projected to two dimensions using t-distributed Stochastic Neighbor Embedding (t-SNE)[3] with correlation dissimilarity (first 50 principle components, or the number of electrodes if lower). (**B**) IR dynamics for the regions and categories presented in (A). Dashed lines: stimulus onset and chance level (no single-item information). Colored vertical lines on the abscissa indicate peak times (plotted in (A)). Statistical significance assessed using cluster permutations (red horizontal bars mark significant clusters). (**C**) IR stability matrices corresponding to time-courses in (B) (dashed line on the diagonal). Significant clusters (cluster-based permutations) are contoured in black; corresponding cluster p-value is shown in the top-right corner. (**D**) IR dynamics for all categories in the four main ROIs. Notations and significance as in (B). Single exemplar information was present in multiple categories in VT and Occ. IR stability matrices (not shown) showed stability for objects and animals in both regions (all $p_{cluster} < 0.006$) and transient representation for faces (both $p_{cluster} < 0.042$). Unsurprisingly, no region showed reliable representation of single watches, as this category was designed to be highly homogeneous (Figure 1A). Par did not show significant exemplar information in single categories (though $Par_{faces}$ was marginal, $p_{cluster} < 0.06$). PFC showed high IR within the faces category. Results using 50ms smoothing were similar, except $Occ_{faces}$ was not significant and $Occ_{watches}$ was significant ($p_{cluster} < 0.017$).

***Supplementary Tables***

| Patient | Gender | Age | Handedness |
|---|---|---|---|
| **S1** | F | 38 | R |
| **S2** | M | 46 | R |
| **S3** | M | 41 | L |
| **S4** | M | 19 | R |
| **S5** | F | 42 | R |
| **S6** | M | 29 | R |
| **S7** | M | 47 | L |
| **S8** | F | 65 | R |
| **S9** | M | 47 | R |
| **S10** | F | 36 | R |

**Table S1. Patent demographic details.**

| **Patient** | **Faces** | **Objects** (non-watch) | **Animals** | **Watches** | **Other** (mostly houses and body parts) | **Targets** (both types) | **Total** |
|---|---|---|---|---|---|---|---|
| **S1** | 208 | 124 | 64 | 208 | 20 | 64 | 688 |
| **S2** | 208 | 124 | 64 | 208 | 20 | 64 | 688 |
| **S3** | 104 | 62 | 32 | 104 | 10 | 32 | 344 |
| **S4** | 102 | 59 | 37 | 103 | 11 | 32 | 344 |
| **S5** | 204 | 118 | 74 | 206 | 22 | 64 | 688 |
| **S6** | 102 | 59 | 37 | 103 | 11 | 32 | 344 |
| **S7** | 203 | 128 | 68 | 205 | 20 | 64 | 688 |
| **S8** | 199 | 127 | 68 | 204 | 20 | 64 | 682 |
| **S9** | 102 | 59 | 37 | 103 | 11 | 32 | 344 |
| **S10** | 102 | 59 | 37 | 103 | 11 | 32 | 344 |

**Table S2. Number of stimuli viewed by each patient (all durations).**

| Patient | Implanted hemisphere | Coverage | Number of electrodes | | |
|---|---|---|---|---|---|
| | | | **All** | **Noise-free** | **Responsive** |
| **S1** | R | Occ, VT, Par, PFC, LT, Med | 64 | 63 | 33 |
| **S2** | R | Occ, VT, Par, SM, LT, Med | 112 | 96 | 53 |
| **S3** | R | Occ, VT, Par, SM, LT, Med | 118 | 107 | 77 |
| **S4** | R | Occ, VT, PFC, Par, SM, LT | 116 | 116 | 22 |
| **S5** | R | Occ, VT, Par, PFC, SM, LT, Med | 100 | 97 | 39 |
| **S6** | R | Occ, VT, Par, PFC, SM, LT, Med, Depth (4 elec) | 98 | 88 | 27 |
| **S7** | L | VT, Par, PFC, LT | 64 | 26 | 16 |
| **S8** | R | Occ, VT, Par, PFC, SM, LT | 110 | 93 | 53 |
| **S9** | L | Occ, VT, Par, SM, LT, Med | 128 | 100 | 51 |
| **S10** | 110 R, 12 L (depth) | Occ, VT, Par, PFC, SM, LT, Depth (24 elec) | 122 | 121 | 59 |

**Table S3. Individual electrode coverage.**
Occ, occipital; VT, ventral-temporal; PFC, prefrontal; Par, parietal; SM, sensorimotor; LT, lateral-temporal; Med, Medial.

| **Region** | **All electrodes** | **Noise-free** | **Responsive** |
|---|---|---|---|
| **Occipital (Occ)** | 174 | 159 | 126 |
| Occipital retinotopic | 94 | 90 | 76 |
| Occipital non-retinotopic | 80 | 69 | 50 |
| **Ventral-temporal (VT)** | 255 | 227 | 161 |
| **Parietal (Par)** | 171 | 138 | 43 |
| **Prefrontal (PFC)** | 108 | 101 | 36 |
| Orbito-frontal cortex (OFC) | 35 | 30 | 14 |
| Lateral prefrontal cortex (LPFC) | 73 | 71 | 22 |
| **Sensorimotor (SM)** | 90 | 89 | 13 |
| **Lateral-temporal (LT)** | 188 | 153 | 31 |
| Medial (not analyzed) | 18 | 12 | 4 |
| Depth (not analyzed) | 28 | 28 | 16 |
| **Total** | 1032 | 907 | 430 |

**Table S4. Number of electrodes per region.**

| Time-window | Faces | Objects (non-watch) | Animals | Watches | Any category |
|---|---|---|---|---|---|
| **100-300ms** | 275 | 219 | 199 | 235 | 342 |
| **300-500ms** | 220 | 225 | 204 | 198 | 347 |
| **500-700ms** | 132 | 133 | 128 | 104 | 219 |
| **700-900ms** | 94 | 78 | 53 | 69 | 142 |
| **Any window** | 313 | 285 | 246 | 281 | 430 |

**Table S5. Number of responsive electrodes per category and time-window.**

| **Patient group** | **Faces** | | | **Objects** (non-watch) | | | **Animals** | | | **Watches** | | |
|---|---|---|---|---|---|---|---|---|---|---|---|---|
| | **300** | **900** | **1500** | **300** | **900** | **1500** | **300** | **900** | **1500** | **300** | **900** | **1500** |
| **S1-S10** | 6 | 8 | 3 | 4 | 1 | 4 | 5 | 3 | 2 | 8 | 6 | 8 |
| **S1-S3** | 20 | 19 | 20 | 10 | 9 | 14 | 6 | 9 | 5 | 20 | 20 | 20 |
| **S4-S10** | 32 | 30 | 26 | 17 | 17 | 18 | 12 | 12 | 9 | 32 | 28 | 30 |

**Table S6. Number of unique exemplars viewed by each decoding patient group.**
Trials containing ictal spikes or excessive noise were excluded.

| **Patient group** | **Occ** | | | **VT** | **Par** | **PFC** | | | **SM** | **LT** |
|---|---|---|---|---|---|---|---|---|---|---|
| | Total | Retin | Non-retin | | | Total | OFC | LPFC | | |
| **S1-S10** | 126 | 76 | 50 | 161 | 43 | 36 | 14 | 22 | 13 | 31 |
| **S1-S3** | 69 | 43 | 26 | 69 | 16 | 0 | 0 | 0 | 0 | 5 |
| **S4-S10** | 57 | 33 | 24 | 92 | 27 | 36 | 14 | 22 | 13 | 26 |
| **S1-2, S5, S7-8** | 37 | 24 | 13 | 88 | 18 | 21 | 9 | 12 | 12 | 14 |
| **S1, S2, S5-S10** | 67 | 41 | 26 | 134 | 33 | 35 | 14 | 21 | 12 | 30 |

**Table S7. Number of responsive electrodes in each patient subgroup.**
